# Human skeletal muscle organoids model fetal myogenesis and sustain uncommitted PAX7 myogenic progenitors

**DOI:** 10.1101/2020.09.14.295832

**Authors:** Lampros Mavrommatis, Hyun-Woo Jeong, Urs Kindler, Gemma Gomez-Giro, Marie-Cécile Kienitz, Martin Stehling, Olympia E. Psathaki, Dagmar Zeuschner, M. Gabriele Bixel, Dong Han, Gabriela Morosan-Puopolo, Daniela Gerovska, Ji Hun Yang, Jeong Beom Kim, Marcos J. Araúzo-Bravo, Jens C. Schwamborn, Stephan A. Hahn, Ralf H. Adams, Hans R. Schöler, Matthias Vorgerd, Beate Brand-Saberi, Holm Zaehres

## Abstract

In vitro culture systems that structurally model human myogenesis and promote PAX7^+^ myogenic progenitor maturation have not been established. Here we report that human skeletal muscle organoids can be differentiated from induced pluripotent stem cell lines to contain paraxial mesoderm and neuromesodermal progenitors and develop into organized structures reassembling neural plate border and dermomyotome. Culture conditions instigate neural lineage arrest and promote fetal hypaxial myogenesis towards limb axial anatomical identity, with generation of sustainable uncommitted PAX7 myogenic progenitors and fibroadipogenic (PDGFRa+) progenitor populations equivalent to those from the second trimester of human gestation. Single cell comparison to human fetal and adult myogenic progenitors reveals distinct molecular signatures for non-dividing myogenic progenitors in activated (CD44^High^/CD98^+^/MYOD1^+^) and dormant (PAX7^High^/FBN1^High^/SPRY1^High^) states. Our approach provides a robust 3D in vitro developmental system for investigating muscle tissue morphogenesis and homeostasis.

## Introduction

Novel skeletal muscle model systems are required to further elucidate the process of human myogenesis as well as investigate muscular disorders and potential gene, cell or drug therapies. Two-dimensional (2D) culture conditions guide pluripotent stem cell (PSC) differentiation towards skeletal muscle lineage using sequential growth factor applications and/or conditional PAX7 expression (Chal et al., 2015, Xi et al., 2017, Shelton et al., 2014, Borchin et al., 2013, Darabi et al., 2012). Further, surface marker expression can be utilized to isolate myogenic progenitors with in vivo repopulation potential (Magli et al., 2017, Hicks et al., 2018, Al Tanoury et al., 2020, Sun et al., 2022). While the few described three-dimensional (3D) differentiation approaches have provided cohorts of terminally differentiated myofibers, focus on potential interactions with the vasculature and nervous system has neglected assessment of the developmental identity or sustainability of myogenic progenitors (Faustino Martins et al., 2020, Maffioletti et al., 2018, Rao et al., 2018). Single cell technologies increasingly provide databases for deciphering myogenic trajectories and expression profiles of myogenic stem and progenitor cells (Barruet et al., 2020, Rubenstein et al., 2020, Xi et al., 2020), enabling full evaluation of the ability of PSC differentiation protocols to mimic human development. Translation to model muscular dystrophies and investigate potential interventions in vitro necessitates methods that provide expandable populations of muscle progenitors while promoting self-renewal and preserving a quiescent, non-dividing, state (Quarta et al., 2016, Montarras et al.,2005).

Most vertebrate skeletal muscle progenitors develop from the paraxial mesoderm via transient embryonic developmental structures (somites and dermomyotome) into the skeletal muscle system that spans the whole body. Here, we evaluate human skeletal muscle organoids as a novel system to structurally investigate myogenic differentiation from human induced pluripotent stem cells (iPSCs) in a 3D environment, mimicking pathways described for chicken and mouse (Buckingham and Rigby, 2014). We develop a comprehensive supplementation/reduction protocol to first drive differentiation towards paraxial mesoderm through application of the GSK3 inhibitor CHIR99021, BMP inhibitor LDN193189 and bFGF. Subsequent stimulation with WNT1A, SHH, FGF and HGF is designed to promote derivation of organized structures reassembling neural plate border and dermomyotome. We then aim to arrest neural lineage via FGF removal, while stimulating with HGF to selectively promote propagation of undifferentiated myogenic progenitors and consequent generation of fetal myofibers. Our goal is to provide PAX7^+^ myogenic progenitors in a non-dividing quiescent state sustainable over weeks after differentiation induction. Single cell analysis will position cells along the quiescent-activation myogenic trajectory discriminating dormant (PAX7+, FBN1+, SPRY11+, CHODL1+) and activated (CD44+, CD98+, MYOD1+, VEGFA+) states. We thus seek to develop and validate a new skeletal muscle organoid system for investigating human myogenesis with translational potential for disease modeling and therapy development.

## Results

### Generation of human fetal skeletal muscle organoids

Organoid cultures from pluripotent stem cells often require pre-patterning before Matrigel embedding to promote structural development (Lancaster et al., 2013; Spence et al., 2011; Koehler et al., 2017). In our 3D approach, we provided cells with immediate matrix support upon embryoid body formation to preserve cell state transitions. Initial pre-embedding screening indicated high expression of pluripotent markers OCT4, SOX2 and NANOG, moderate expression of neural tube marker PAX6 and low expression of mesodermal markers BRACHYURY and MSGN1 **(Figures 1, 1-figure supplement 1A)**. Upon Matrigel embedding, stimulation with CHIR99021, LDN193189 and bFGF promoted paraxial mesoderm formation through derivation of BRACHYURY^+^ and TBX6^+^ cells **(Figures 1A,C, 1-figure supplement 1A)**. Immunostaining at Day 5 depicted presence of mesodermal (BRACHYURY^+^), paraxial mesodermal (TBX6^+^) and neuromesodermal (SOX2^+^/BRACHYURY^+^) progenitors (Gouti et al., 2017; Henrique et al., 2015) **(Figure 1C).** From this stage, we attempted to mimic determination front formation and promote anterior somitic mesoderm (ASM) (Aulehla and Pourquie,2010; Shimozono et al., 2013) by maintaining constant CHIR99021 and LDN193189 levels, reducing bFGF levels to 50% and simultaneously introducing retinoic acid to the culture. Consequently, distinct PAX3^+^ but SOX2^-^ cells emerged on the organoid surface **(Figure 1D)**, followed by a significant downregulation of PSM markers HES7, TBX6 and MSGN1 **(Figures 1E, 1-figure supplement 1A)**. Concomitantly, we observed significant upregulation of ASM marker MEOX2 and neural crest marker TFAP2A but not of SOX2 or PAX6, excluding a shift towards neural tube formation **(Figures 1E,F, 1-figure supplement 1A)**. Up to this stage, inter-organoid sizes showed small variability **(Figure 1B)**. Dermomyotomal fate was promoted by Sonic Hedgehog (SHH) and WNT1A stimulation, while maintaining BMP inhibition avoided lateral mesoderm formation **(Figure 1A)**. Expression profiling at Day 11 depicted presence of neural tube/crest (SOX2, PAX6, TFAP2A, SOX10 upregulation) and mesodermal (UNCX, TBX18, PAX3) lineages and further downregulation of PSM markers **(Figures 1E,F, 1-figure supplement 1A)**. Notably, the dermomyotomal/neural crest marker PAX7 emerged together with markers that define the dorsomedial (EN1) or ventrolateral portion (SIM1) of the dermomyotome (Cheng et al., 2004) **(Figures 1D,F, 1-figure supplement 1A)**.

**Figure 1.**
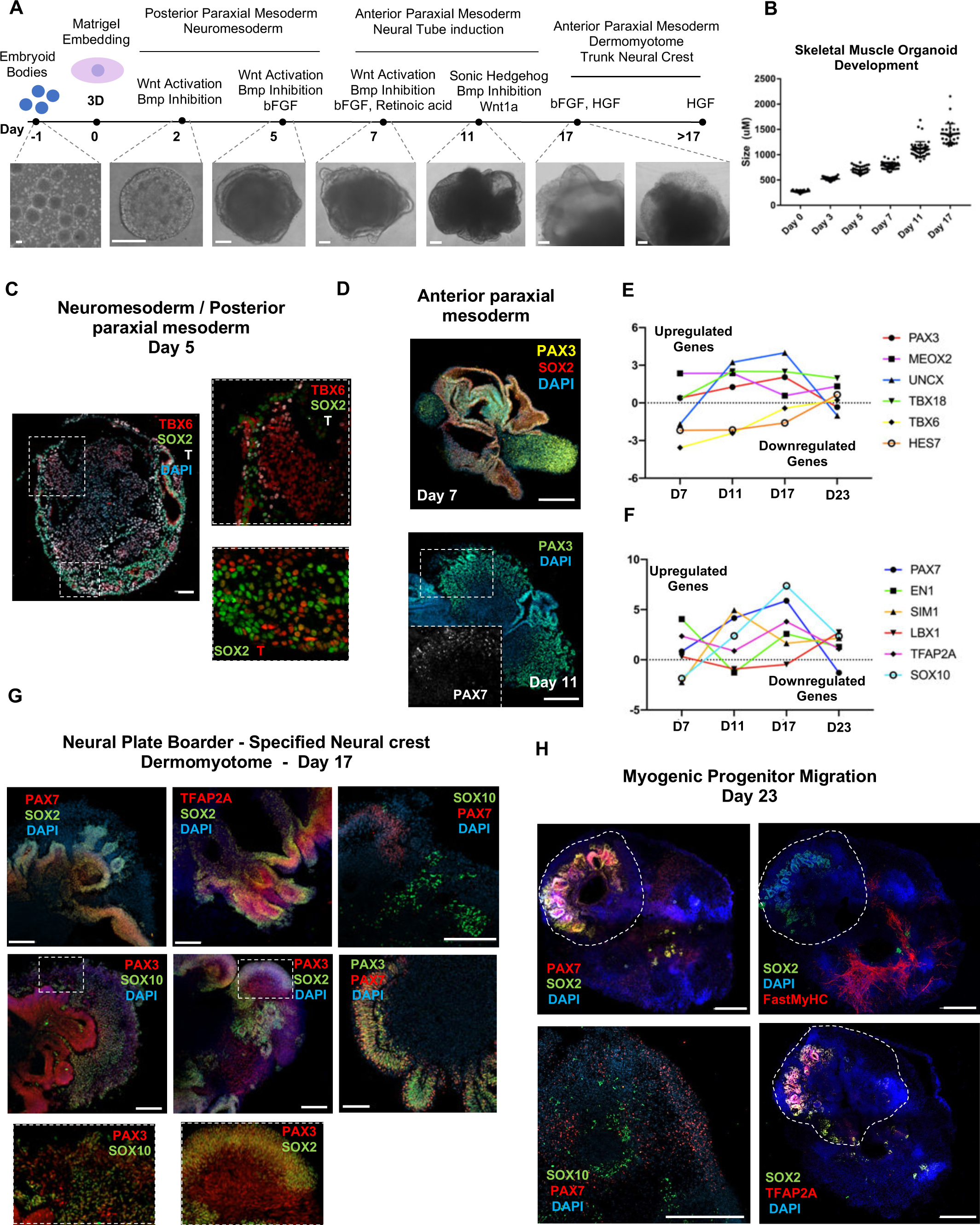
Skeletal muscle organoid protocol and correlation to fetal development. **(A)** Brightfield images of myogenic development stages, with corresponding cytokines/growth factors protocol applications **(B)** Graph depicting organoid development in size (Day 0, *n=* 51; Day 3, *n=* 46; Day 5, *n=* 48; Day 7, *n=* 50; Day 11, *n=* 50; Day 17, *n=* 33; for each timepoint organoids from 3 independent derivations were measured) **(C)** Representative mesodermal T+, neuromesodermal T+, SOX2+ and paraxial mesodermal TBX6+ organoid expression (in live DAPI-cells) at Day 5 **(D)** Representative organoid PAX3+, mesodermal SOX2- and neural SOX2+ expression at Day 7; PAX3/PAX7 coexpression at Day 11 **(E)** Graph depicting qPCR values for anterior PAX3/UNCX/MEOX2/TBX18 and posterior TBX6/HES7 somitic mesodermal markers **(F)** Graph depicting qPCR values for epaxial and hypaxial dermomyotomal PAX7/EN1/SIM1/LBX1 and neural crest TFAP2A/SOX10 markers **(G)** Representative organoid neural plate border epithelial PAX3+ / PAX7+ / SOX2+ / TFAP2A+, paraxial mesodermal PAX3+/SOX2- and delaminating specified neural crest progenitor PAX3+ / SOX10+ expression at Day 17 **(H)** Representative organoid myogenic FastMyHC+, PAX7+ and neural SOX2+/TFAP2A+/SOX10+ expression at Day 23. Dashed line represents location of embryoid body embedded into Matrigel. Statistics: Values at each timepoint represent difference in mean relative expression for each gene (D7= Day 5 - Day 7, D11= Day 7 - Day 11, D17= Day 11 - Day 17, D23= Day 17 - Day 23) as derived by performing ordinary one-way ANOVA and Tukey ′s multiple comparison tests (E,F). Scale bars: 200μm (G), 100μm (A,D,H), 50μm

Consequently, for favoring myogenesis we stimulated organoid culture with FGF and HGF (Charge and Rudnicki, 2004) **(Figure 1A)**. Surprisingly, at Day 17, organoids constituted a mosaic of neural crest and myogenic progenitor cells. Cells with epithelial morphology were TFAP2A^+^, SOX2^+^, PAX3^+^ and PAX7^+^, indicating formation of neural plate border epithelium (Roellig et al., 2017) **(Figure 1G)**. In cell clusters with mesenchymal morphology, we detected specified migrating PAX3^+^/SOX10^+^ neural crest progenitors and PAX3^+^ /SOX2^-^/ SOX10^-^ cells of myogenic origin **(Figure 1G)**. At this stage, myogenic lineages appeared to be primarily represented by PAX3^+^ (9.35±0.07%) rather than PAX7^+^ cells, as the PAX7 expression pattern (15.3±0.1% PAX3^+^/PAX7^+^, 5.11±0.13% PAX7^+^) predominantly overlapped with that of SOX2 and TFAP2A **(Figures 1G, 1-figure supplement 1B)**. Morphologically, neural plate boarder and dermomyotomal populations exhibited uneven distribution, and thereby subsequent neural crest and myogenic migration processes introduced organoid size variability **(Figure 1B,G)**. From Day 17 onwards, we omitted FGF stimulation to cease neural crest development and promote delamination/migration of PAX3^+^/ PAX7^+^/SOX10^-^ progenitor cells (Murphy et al., 1994) **(Figure 2A)**. Strikingly, until Day 23, we observed committed myogenic populations through detection of fast MyHC myofibers in proximity of PAX7^+^ but SOX2^-^/TFAP2A^-^/SOX10^-^ negative cells **(Figures 1H, 1-figure supplement 1C to E)**. Consistently, expression profiling indicated downregulation of neural tube/crest lineage markers and significant upregulation of muscle precursor migrating markers (Buckingham and Rigby, 2014), such as LBX1, CXCR4 and MEOX2 **(Figures 1E, 2A, 2-figure supplement 2A)**.

**Figure 2.**
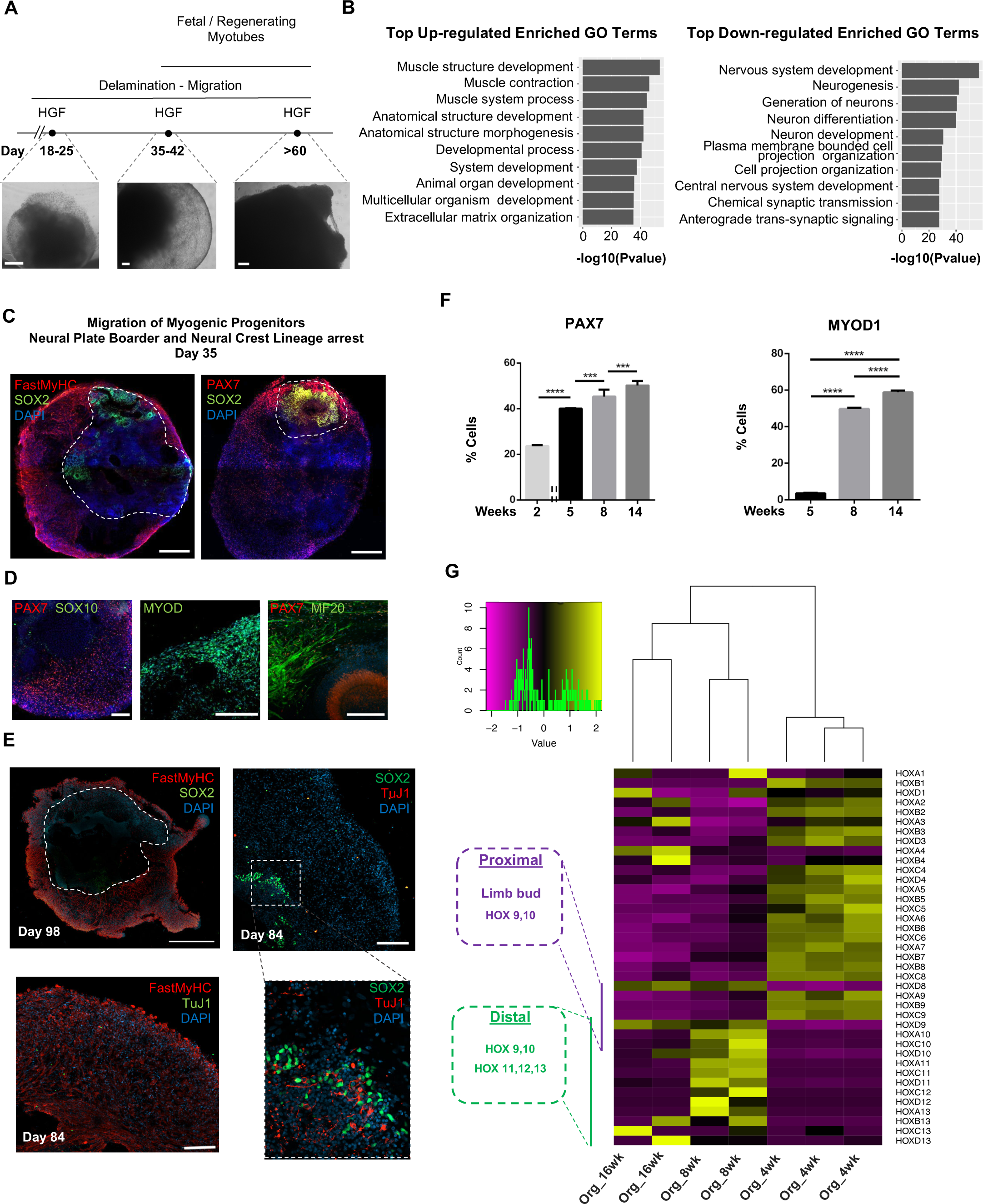
Neural lineage development during skeletal muscle organoid progression. **(A)** Stepwise brightfield images depicting delamination / migration of progenitor population during organoid culture progression and corresponding myofiber formation **(B)** Gene ontology enrichment analysis comparing 4 and 8 weeks organoids attributes muscle identity at 8 weeks post differentiation and highlights muscle system development and neural lineage arrest among the top upregulated and downregulated gene ontology terms, respectively **(C-E)** Organoid overview at Day 35 indicates predominant expression of FastMyHC^+^ and PAX7^+^ myogenic populations, while SOX2^+^ neural populations demarcate SOX2 neural plate border epithelium location as observed at earlier stages (Day 16) **(C)**; PAX7 cells are of myogenic origin (PAX7^+^/SOX10^-^), MF20^+^ myotubes are in their proximity and MYOD1^+^ cells appear at organoid periphery **(D)**; TUJ1^+^ neurons are restricted to inner organoid areas and close to SOX2^+^ epithelium, while FastMyHC^+^ myofibers occupy exterior organoid areas**(E) (F)** Histographs based on FACS intracellular quantification depicting percentage of PAX7^+^ or MYOD1^+^ cells through differentiation protocol. For each replicate 10 organoids were pooled, n=10 Statistics: *P < 0.05, **P < 0.01, ***P < 0.001, ****P < 0.0001, ns: not significant **(G)** Heatmap of HOX gene cluster emphasizes organoid culture limb axial anatomical identity, by depicting transition from an initial limb bud (HOX 9-10) towards a more distal identity (HOX 11-13) at 8 weeks and 16 weeks post differentiation, respectively. Scale bars, 500μm (C), 200μm (A, D, E)

At 8 weeks, organoids showed profound changes in transcription profiling **(Figures 2, 2-figure supplement 2B).** Gene ontology enrichment analysis highlighted an ongoing developmental transition with muscle development, muscle contraction and myofibril assembly among the top upregulated gene ontology terms and neurogenesis and nervous system development among the top downregulated terms **(Figure 2B)**. In addition, we detected downregulation of key markers characterizing neural tube and neural crest development (Soldatov et al., 2019), such as PAK3, DLX5, B3GAT1, FOXB1, HES5 and OLIG3 **(Figures 2, 2-figure supplement 2C)**. Interestingly, we could spatially visualize this process using immunocytochemistry for SOX2, TFAP2A and SOX10 expressing cells that were restricted to the inner portion of the organoid, and probably not susceptible to culture stimulation at 5 weeks **(Figures 1H, 2C,D, 2-figure supplement 2D)**. This neural / myogenic lineage spatial orientation could be visualized even at Day 84 through presence of TUJ1^+^ neurons confined to inner organoid areas and close to SOX2^+^ epithelium, while FastMyHC^+^ myofibers occupied exterior areas **(Figure 2E).** On the other hand, substantial proportions of migratory cells in proximity of FastMyHC^+^, MF20^+^ myofibers expressed PAX7 (40±0.3%) but not SOX2, TFAP2A, SOX10 or MYOD1 (3.55±0.32%) (**Figures 2F, 2-figure supplement 2D,E**). This behavior is further illustrated by the presence of MYOD1^+^ cells confined to organoid periphery **(Figure 2D)**.

Culture progression led to significant increase in PAX7^+^ and MYOD1^+^ cells at 8 weeks (45.3±3,4% and 49.8±0.62%, respectively) (**Figures 2F, 2-figure supplement 2E)**, suggesting an interval in which organoid culture predominantly commits to myogenic lineage. Expression profiling at this stage depicted upregulation of markers characteristic for limb migrating myogenic cells, e.g. LBX1, PAX7, SIX1/4, EYA1/4, PITX2, MYF5, TCF4 and MSX1 **(Figures 2, 2-figure supplement 2A)**. Consequently, HOX gene cluster profiling at 4 weeks correlated HOX A9, B9, C9 upregulation with limb bud site, while upregulation of the HOX 10-13 cluster during 8 to16 weeks attributed later organoid development to a more distal limb axial anatomical identity (Shubin et al., 1997; Xu et al., 2011, Raines et al., 2015) **(Figure 2G)**.

### Lineage representation and developmental identity for skeletal muscle organoids

Analysis of organoid culture at single cell resolution indicated predominant presence of skeletal muscle lineage (n= 3945 cells, 91.26% of total population), complemented with two smaller cell clusters of mesenchymal / ‘fibroadipogenic’ (n=165 cells, 3.82% of total population), and neural (n=213 cells, 4.93% of total population) origin **(Figures 3A, 3-figure supplement 1A**). Skeletal muscle lineages further separated into distinct sub-clusters: myogenic muscle progenitors in non-dividing (PAX7^+^/PAX3^-^) and mitotic states (PAX7^+^/CDK1^+^/KI67^+^), myoblasts (MYOD1^+^), myocytes (MYOG^+^) and myotubes (MYH3^+^) **(Figures 3A, 3-figure supplement 1A,B)**. Further, investigation of mesenchymal cluster identity suggested fibroadipogenic potential of a distinct cell population with PDGFRa among the top upregulated genes (Uezumi et al., 2010,2011; Xi et al., 2020) **(Figures 3A,F, 3-figure supplement 1A,B).** Consistently, we detected upregulation of fibrotic markers. Moreover, adipogenic potential was highlighted through upregulation of PREF-1 and EBF2 **(Figure 3F)**. Occasionally, we detected structures resembling adipocytes from 9 to10 weeks onwards, with adipogenic nature verified by PRDM16 and Oil red O positive staining **(Figures 3B,D, 3-figure supplement 1C,D)**. A SOX2^+^/TUBB3^+^ positive neural cluster potentially derived from the neural plate border epithelium at younger stages could be located towards organoid interior **(Figures 3C, 3-figure supplement 1E).**

**Figure 3.**
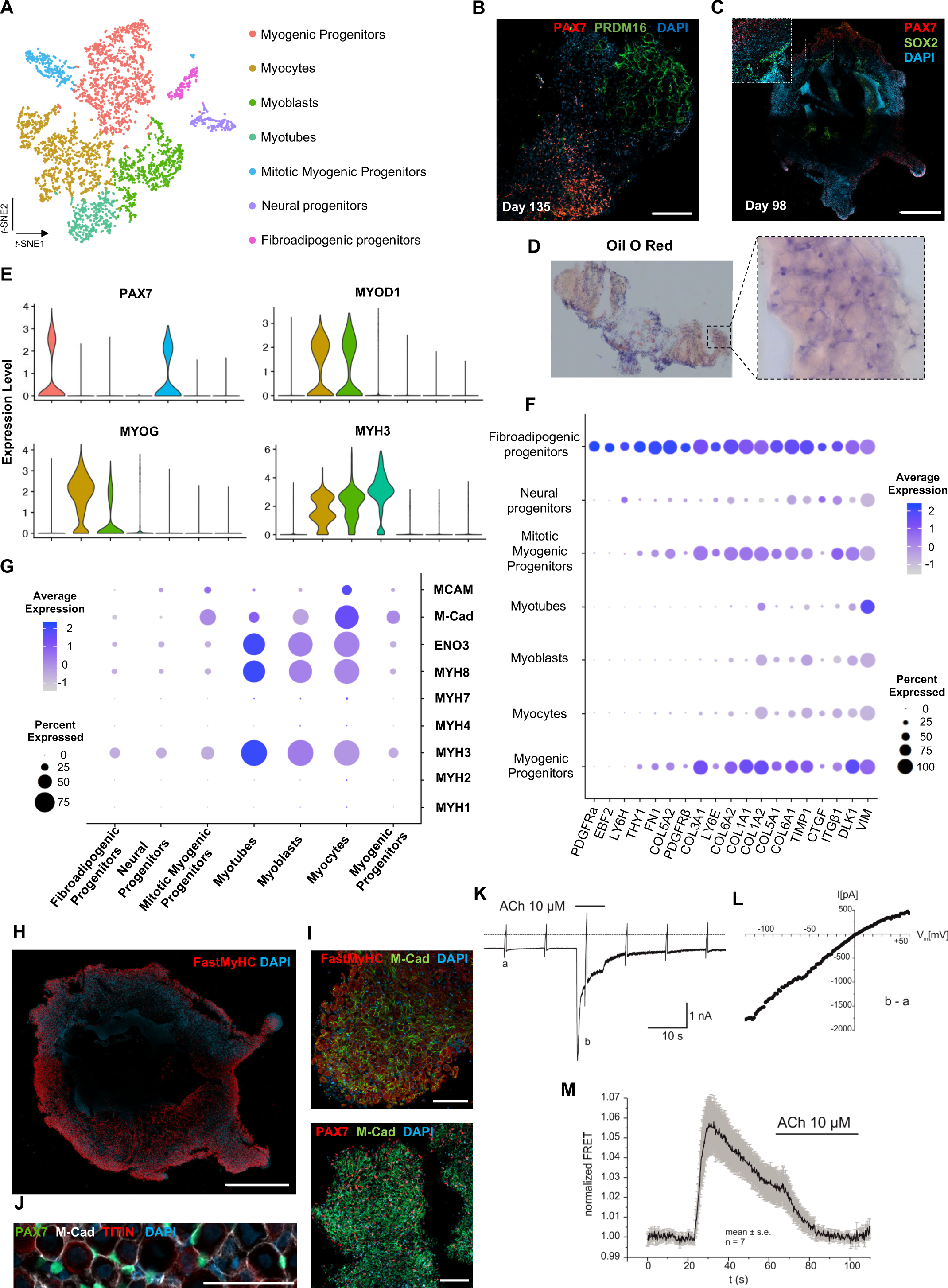
Skeletal muscle organoid characterization at single cell resolution. **(A)** t-SNE visualization of color-coded clustering (n= 4.323 cells) at 12 weeks post differentiation highlights predominant presence of skeletal muscle lineage, represented by clusters corresponding to myogenic progenitors (n=1625 cells, 37% of total population) in non-dividing (n=1317 cells) and mitotic (n=308 cells) state, myoblasts (n=731 cells), myocytes (n=1147 cells) and myotubes (n=442). Additionally, mesenchymal and neural lineages are represented by 2 smaller clusters of fibroadipogenic (n=165 cells) and neural (n=213 cells) progenitors, respectively **(B)** Immunocytochemistry at Day 135 indicates derivation of PRDM16^+^ adipocyte clusters distinct from PAX7^+^ myogenic progenitors **(C)** Organoid overview at Day 98 depicts expression of PAX7^+^ myogenic populations and highlights SOX2+ neural populations towards organoid interior **(D)** Positive areas with Oil O Red staining indicate derivation of adipocytes in organoid culture at Day 135 **(E)** Violin plots of key markers PAX7, MYOD1, MYOG, MYH3 from each stage as in 3A depict relative expression levels and emphasize gradual transition from myogenic progenitor to myotube subcluster **(F)** Dot plot showing expression of representative genes related to adipogenesis and fibrogenesis across the 7 main clusters. Circle area represents percentage of gene^+^ cells in a cluster, color reflects average expression level (gray, low expression; blue, high expression) **(G)** Dot plot showing expression of representative genes related to fetal myogenesis across the 7 main clusters. Circle area represents percentage of gene^+^ cells in a cluster, color reflects average expression level (gray, low expression; blue, high expression). **(H-J)** Representative organoid overview at Day 98 indicates predominant expression of Fast MyHC fetal myofibers **(H,J),** positive for M-Cadherin and in PAX7^+^ cells proximity **(I,J) (K)** Representative recording (n=6) of acetylcholine (ACh)-induced changes in holding current in a single skeletal muscle cell. ACh (10 μM) was applied as indicated by the bar. Holding potential -90 mV. Downward deflections represent membrane currents in response to depolarizing voltage ramps (duration 500 ms) from -120 mV to +60 mV. Dashed line indicates zero current level. **(L)** I/V curve of nAChR currents obtained by subtraction of voltage ramp-induced changes of current in presence and absence of ACh (10-5 mol/L), corresponding lower-case letters b - a **(M)** Summarized FRET-recordings from skeletal muscle cells transfected with Twitch2B to monitor the increase in [Ca2+]i during ACh application (h) Scale bars, 1mm (C,H), 200μm (B), 100μm (I,J)

Myogenesis progression based on *t*-SNE feature and Violin plots of key markers from each stage indicated gradual transitions from myogenic progenitor to myotube subclusters (**Figures 3E, 3-figure supplement 1F**). MyHC expression profiling at single cell resolution demonstrated predominant presence of embryonic (MYH3^+^) and perinatal (MYH8^+^) myofibers, co-expressing β-Enolase and M-cadherin in vicinity of PAX7^+^ cells (**Figure 3** G - J). Consistently, we detected MCAM expression on muscle subclusters (**Figure 3G**), while bulk RNA seq at 16 weeks highlighted expression of additional MyHC isoforms, e.g. MYH2 (II_a_), MYH4(II_b_) and transcription factor NFIX (Moore et al., 1993; Barbieri et al., 1990; Alexander et al. 2016; Messina et al., 2010). At more mature stages, we detected adult MYH1 or slow MYH7 isoforms (**Figures 3, 3-figure supplement 1G**), presumably due to myofiber stimulation via spontaneous contraction. In agreement, differential expression comparison between 8 and 16 week organoids indicated less variance in transcription profiling (**Figures 3, 3-figure supplement 1H, 4A**). Consistently, sarcomere organization, ion transport, response to stimulus and synapse structure maintenance were among the upregulated gene ontology terms and mitosis, cell cycle, DNA packaging and nuclear division among the top downregulated terms (**Figures 3, 3-figure supplement 2B-D**). Notably, ongoing maturation did not affect pool of progenitor cells, as even at 14 weeks we could report significant increases in PAX7^+^ (50,16±2.19%) and MYOD1^+^ (58.53±0.92%) cells (**Figures 2F, 3, 3-figure supplement 2E,F)**.

### Functionality and maturation of organoid derived myofibers

Regarding localization and functionality of organoid derived myofibers, immunocytochemistry revealed positive staining for dystrophin and a continuous laminin sheath around individual muscle fibers. Ultrastructure and 2-photon microscopy analysis depicted well developed sarcomeres **(Figures 3, 3-figure supplement 3A-D)**. Moreover, patch clamp experiments, upon superfusion of cells with acetylcholine (ACh), indicated inward current generations that rapidly declined to a low steady state level **(Figure 3K, 3-figure supplement 3E**). The I/V-curve of this current component showed almost linear behavior and reversal potential around 0 mV **(Figure 3L)**. These data are in line with previous studies that described currents with analogous properties as characteristic nAChR currents (Jahn et al., 2001; Shao et al., 1998). Application of a fluorescent biosensor for monitoring changes in cytosolic Ca^2+^ (Twitch2B) revealed nAChR efficiency in modulating intracellular [Ca^2+^]. Summarized FRET recordings **(Figures 3M, 3-figure supplement 3F)**, following application of ACh (10 μM), illustrated a rapid increase in FRET ratio that gradually declined in the presence of ACh, probably reflecting desensitization of nAChRs. These results demonstrated that nAChR in skeletal muscle cells are functional in terms of inducing Ca^2+^ release from intracellular stores.

Using our organoid protocol, we successfully derived fetal muscle progenitors and electrophysiologically functional myofibers from hiPSC lines with wild type and Duchenne Muscular Dystrophy genetic backgrounds (**Figures 3, 3-figure supplement 3E,F**).

### Identity and sustainability of organoid derived PAX7 myogenic progenitors

Skeletal muscle organoids remarkably foster sustainable propagation of PAX7^+^ progenitors **(Figures 2F, 3A)**. Initial screening of PAX7^+^ progenitors indicated expression of several satellite cell markers (Fukada et al., 2007), such as CD82, CAV1, FGFR1, FGFR4, EGFR, M-Cadherin, NCAM, FZD7, CXCR4, ITGβ1, ITGA7, SCD2 and SCD4 **(Figures 4, 4-figure supplement 1A)**. Further investigation verified NOTCH signaling activity (HES1, HEYL, HEY1 and NRARP) in the myogenic subcluster, while dormant myogenic progenitors exhibited high expression of SPRY1 and cell cycle inhibitors p57 (CDKN1C), p21 and PMP22 (Xi et al., 2020; Fukada et al., 2007; Bjornson et al., 2012; Shea et al., 2010) **(Figures 4, 4-figure supplement 1B, 2A,D)**. Moreover, proliferation assays performed at the same stage (14 weeks) demonstrated 4.78±0.28% edU^+^ cells, while substantial proportions of PAX7^+^ myogenic progenitors remained quiescent **(Figures 4, 4-figure supplement 1C,D)**. t-SNE clustering divided myogenic progenitors into three distinct groups with unique molecular signatures: “CD44^high^”, “FBN1^high^” and “CDK1^+^” clusters **(Figures 4A, 4-figure supplement 2A)**. The “CD44^high^” cluster, further characterized by CD98 upregulation, adopted a molecular signature similar to activated satellite cell state (Porpiglia et al., 2017) **(Figures 4A,B, 4-figure supplement 2A,D).** In this state, myogenic progenitors expressed VEGFA **(4-figure supplement 2A,D)** as previously described for murine satellite cells (Verma et al., 2018). Consistently, the CD44^+^ / PAX7^+^ ‘activated’ myogenic population located at sites more accessible to HGF signaling, e.g. exterior organoid areas and forming bulges **(Figure 4B)**. The “FBN1^High^” sub-cluster was further characterized by “PAX7^high^ / SPRY1^high^/CHODL^high^ / FBN1^high^” expression **(Figures 4A, 4-figure supplement 2A,D).** PAX7^high^ cells within the dormant state co-expressed NOTCH3, JAG1 and LNFG, together with CHODL markers. Further, we verified presence of FBN1^+^ microfibrils, which, compared to CD44^+^ cells, occupied areas without direct access to activation signals **(Figure 4C)**. Lastly, the “CDK1^+^” cluster was the only proliferative population, further expressing markers mediating satellite cell activation and proliferation, such as DEK and EZH2 (Cheung et al., 2012; Juan et al., 2011) **(Figures 4A,D, 4-figure supplement 1A, 2A,B).**

**Figure 4.**
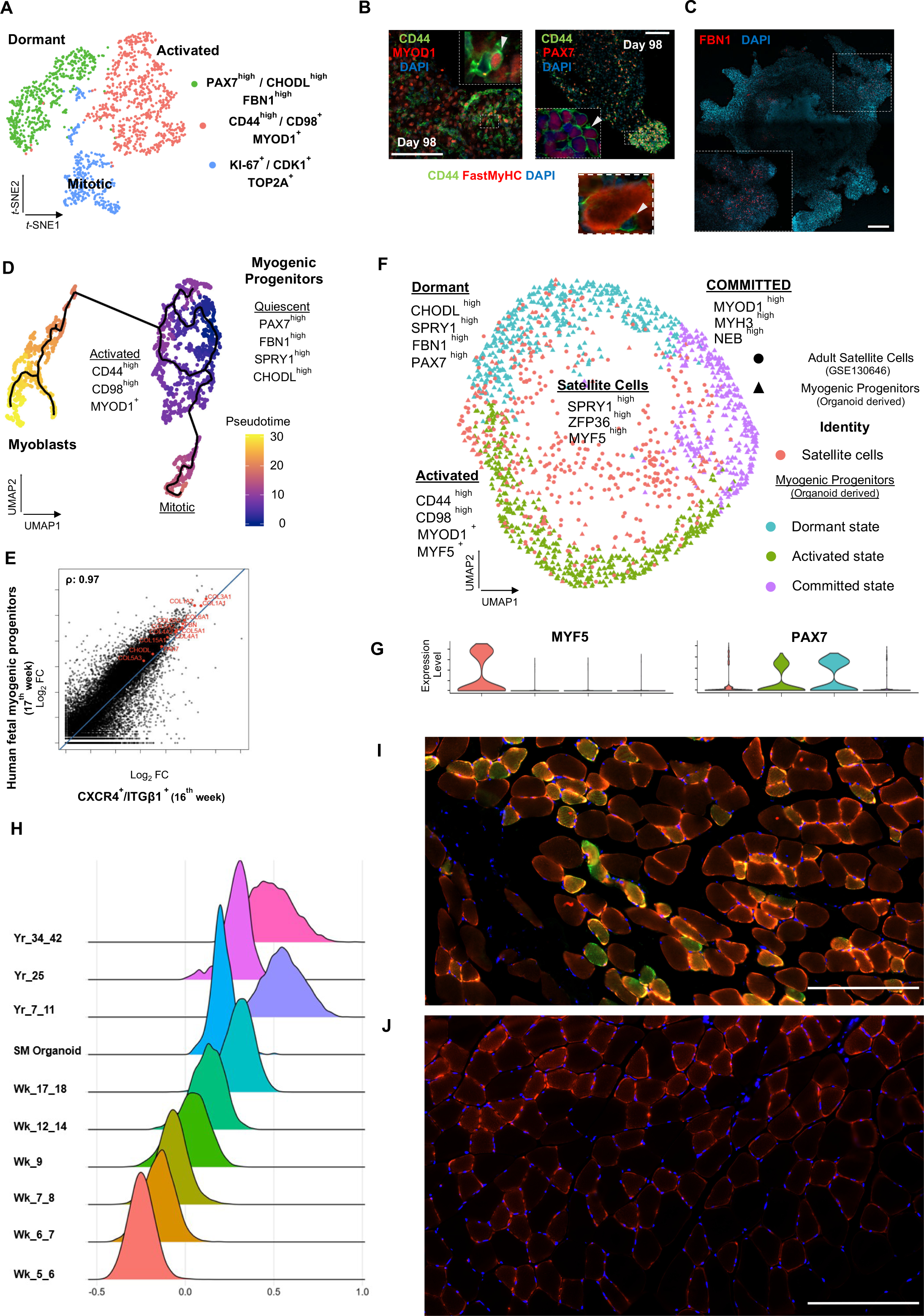
Myogenic progenitor identity and comparison to progenitors derived from fetal and adult muscle tissue. **(A)** t-SNE plot visualization of color-coded clustering indicates myogenic progenitor sub-cluster with distinct molecular signatures: “dormant” PAX7^high^/ CHODL^high^/FBN1^high^, “activated” CD44^high^ / CD98^+^ / MYOD1^+^ and “mitotic” KI-67^+^/CDK1^+^/TOP2A **(B)** Organoid overviews at Day 98 illustrate CD44 and PAX7 expressing cells at developing sites, which are more accessible to HGF activation signal, specificity of CD44 on MYOD^+^/PAX7^+^ progenitor expressing cells (arrows) and absence from FastMyHC^+^ positive myofibers is highlighted **(C)** FBN1^+^ microfibrils are located towards organoid interior **(D)** Pseudotime ordering for myogenic progenitors and myoblast corresponding clusters highlights distinct developmental trajectories promoting myogenic commitment and self-renewal Correlation coefficient plot for Log_2_ fold change (Log_2_ FC) values for isolated myogenic progenitors from human fetal tissue (17w) and FACS sorted CXCR4^+^ / ITGβ1^+^ organoid derived myogenic progenitors (16w). PAX7, COL1A1, COL3A1, COL4A1, COL5A1, COL15A1, FBN1 and CHODL and further extracellular matrix related genes are highlighted on the plot. Pearson’s correlation coefficient, rho=0.9 for global expression comparison and rho=0.97 for selected genes **(E)** UMAP color-based clustering divides non-dividing myogenic progenitors and adult satellite cells into 4 clusters with distinct molecular signatures: satellite cells characterized by SPRY1^high^/ZFP36^high^ /MYF5^high^ expression, co-clustered with dormant SPRY1^high^ / FBN1^high^ / CHODL^high^ / PAX7^high^, activated CD44^high^ / CD98^high^ / MYOD1^+^ and committed, NEB ^high^/ MYH3^high^ / MYOD1^high^ organoid derived myogenic progenitors. Dots correspond to adult satellite cells from GSE130646 database, triangles correspond to organoid derived myogenic progenitors. **(F)** Violin plots depicting relative expression levels of markers specific for quiescent PAX7 or activated MYF5 muscle stem cell state across adult satellite cells (GSE130646) and organoid derived myogenic progenitors subclusters **(H)** Ridge plots of developmental score distribution of myogenic progenitors across in vivo or in vitro stages based on the difference between up-regulated satellite cell and embryonic markers from human reference atlases for weeks (Wk) 5 to 18 embryonic and fetal stages, years (Yr) 7 to 42 adult satellite cells and SM (skeletal muscle) organoid **(I,J)** In vivo engraftment potential of human myogenic progenitors. 100.000 CD82+ sorted human cells were injected into *Tibialis anterior* muscle of nude mice (**I**) Control mice were not injected (**J**). Six weeks post transplantation, transverse cryosections of muscle fibers were stained with huLamin A/C (green), Dystrophin (red) and DAPI (blue). Human cells appear green and red in contrast to murine cells which only show a dystrophin positive red staining. Scale bars 200 μm in I,J

Pseudotemporal ordering of myogenic progenitors indicated that the “FBN1^High^” subcluster was the main progenitor population residing in dormant state, which by downregulating PAX7, FBN1 and CHODL and upregulating CD44, MYOD1, CD98 (SLC3A2) and VEGF generated activated state (**Figures 4D**, 4-figure supplement 2B). In activated state, myogenic progenitors that upregulate MYOD1 entered mitosis, proliferated and followed a trajectory leading to myogenic commitment (**Figures 4D**, 4-figure supplement 2B). This commitment was further accompanied by PARD3, p38a MAPK and CD9 expression (Porpiglia et al., 2017; Troy et al., 2012) (4-figure supplement 2E). Conversely, myogenic progenitors that upregulated NOTCH3, SPRY1 and PAX7 followed a loop trajectory that reinstated dormant stem cell state (4-figure supplement 2D,E). Interestingly, the “FBN1^high^” cluster highly expressed extracellular matrix collagens, e.g. COL4A1, COL4A2, COL5A1, COL5A2, COL5A3 and COL15A1. Notably, such expression declined upon commitment and differentiation (4-figure supplement 2C,F).

### Organoid derived myogenic progenitor comparison to human fetal and adult progenitor/stem cells

To evaluate developmental identity of the myogenic cluster, we isolated ITGβ1^+^ / CXCR4^+^ organoid derived myogenic progenitors via FACS (Garcia et al., 2018) and compared to human fetal muscle progenitors, demonstrating high similarity (Pearson ′s correlation co-efficiency, rho=0.9), with myogenic markers such as PAX7, MYF5, MYOG and MYOD1 at comparable expression levels **(4-figure supplement 3A-F)**. Differential expression comparison verified expression of extracellular matrix collagens and proteins, such as COL4A1, COL5A1, COL6A1, th COL15A1, FBN1 and CHODL, in myogenic progenitors similar to 17^th^ week human fetal tissue (Pearson ′s correlation co-efficiency, rho=0.97) **(Figure 4E)**. Further, to evaluate myogenic potency in vitro, isolated ITGβ1^+^ / CXCR4^+^ organoid derived myogenic progenitor cells were re-plated and allowed to differentiate under the same organoid culture protocol, which demonstrated capacity to generate FastMyHC+ and PAX7+ populations within 10 days **(4-figure supplement 3B,C**). Subsequently, comparison to available transcriptomic dataset of human adult satellite cells at single cell resolution, divided myogenic progenitors and adult satellite cells into four clusters with distinct molecular signatures **(Figure 4F)**. Interestingly, myogenic progenitors were enriched for extracellular matrix proteins, while satellite cells mainly upregulated genes repressing transcription/translation, such as ZFP36 and TSC22D1, or related to early activation response, such as MYF5, JUN and FOS **(4-figure supplement 4A,B).** In line, organoid derived myogenic progenitors exhibited higher NOTCH signaling activity in comparison to satellite cells, with NOTCH3 and RBPJ being among enriched markers **(4-figure supplement 4B**). In contrast, adult satellite cells exhibited PAX7^low^ / MYF5^high^ expression profiling, presumably due to tissue dissociation, thereby indicating a tendency for activation rather than preservation or reinstatement of quiescency (Machado et al., 2017; Seale et al., 2000) **(Figure 4G).** Pseudotime ordering showed two distinct clusters, with adult satellite cells downstream from non-dividing myogenic progenitors **(4-figure supplement 4C).** Consistently, downregulation of genes promoting quiescence like PAX7, NOTCH3 and RBP, and upregulation of activation genes like MYF5, JUN and FOS along the trajectory (**4-figure supplement 4D)** was a further indication that organoid derived myogenic progenitors resided in dormant non-dividing state and that our organoid platform promoted sustainability of myogenic progenitors.

Cell-cell communication analysis of organoids at 12 weeks indicates that myogenic progenitors influence their own fate, mainly with extracellular matrix (ECM) related signals, as well as receive signals predominantly from the mesenchyme but not the neural progenitor cluster (**Figure 4 –figure supplement 5**).

To evaluate for reproducibility of organoid development we applied diffusion map analysis on qPCR-based expression analysis of 32 genes at early stages and integrative analysis on scRNAseq data sets of mature stages of organoid development. The data indicate highly conserved cluster representation of myogenic progenitors at all states, together with skeletal muscle myofibers, fibroadipogenic progenitors and neural progenitors related clusters (**Figure 4 – figure supplement 6**).

Organoid derived myogenic progenitors exhibited expression profiling similar to that of Stage 4 myogenic progenitors, correlating to 17-18^th^ week human fetal development, as reported in the human skeletal muscle atlas (Xi et al., 2020). SPRY1, PLAGL1, POU4F1, TCF4, PAX7, CD82 and CD44 markers, as well as ECM proteins, nuclear factor I family members (NFIA, NFIB, NFIC) and NOTCH signaling are specifically upregulated markers. Furthermore, in comparison to 2D culture protocols, our organoid approach preserved myogenic progenitor dormancy without activating MYOD1 and exhibited higher expression of ECM proteins and key markers characterizing Stage 4 fetal myogenic progenitors (**4-figure supplement 7**). ERBB3 and NGFR markers (Hicks et al., 2017) demarcate populations of earlier developmental stages and were not upregulated either in organoid derived or Stage 4 fetal myogenic progenitors (**4-figure supplement 7**). In addition, organoid myogenic progenitors and Stage 4 fetal myogenic progenitors both downregulated cycling genes like MKI67, TOP2A, EZH2 and PCNA (**4-figure supplement 7**).

In addition, we have performed a developmental score distribution of myogenic progenitors based on the difference between up-regulated satellite cell and embryonic markers from the human reference myogenic atlases (Xi et al., 2020) and adult satellite cell data (Rubenstein et al., 2020, Xi et al., 2020) in comparison to our organoid protocol (**Figure 4H**). Organoid-derived myogenic progenitors represent the late fetal stage of maturation partially overlapping with adult satellite cell scoring. We note the heterogeneity of adult satellite cell populations when performing developmental scoring in line with recent reports (Barruet et al., 2020).

Finally, we transplanted CD82-positive progenitors from our organoids into the *Tibialis anterior* muscle of immunodeficient mice to complement our study with an in vivo experiment (Alexander et al., 2016, Marg et al., 2019, Al Tanoury et al., 2020). The CD82-positivity used for FACS selection of myogenic progenitors prior to transplantation almost exclusively overlaps with Pax7-positive cells, being a subcluster of them (**4-figure supplement 4E**). Six weeks post transplantation we could verify clusters of human Lamin A/C positive cells in the transplanted but not in the control group (**Figure 4I,J**). We have measured the size of myotubes with 41+/-6 µm for the human and 63+/-7 µm for the mice mean diameters (n=15 each).

## Discussion

Human skeletal muscle organoids offer a new cell culture system to study human myogenesis, in particular fetal myogenic progenitors. We demonstrate that modulation of Matrigel embedded embryonic bodies with WNT, BMP and FGF signaling at early stages leads to paraxial mesoderm formation **(Figure 1B)**. Further, under guided differentiation, we could promote concomitant development of neural and paraxial mesodermal lineages and derive mesodermal populations with somitic and dermomyotomal origin **(Figure 1C -F**). With WNT1A and SHH stimulation, neural lineage is directed towards dorsal neural tube / crest development which benchmarks the structural recapitulation of neural plate border epithelium **(Figure 1G)**. In vitro neural plate border can be utilized to assess transcriptomic networks and cell fate decision during human neural crest formation.

Delaminating from the dermomyotome, undifferentiated PAX3 progenitor cells reorientate and align in the cranio-caudal axis to form the underlying myotome or completely detach from the dermomyotome and migrate into the surrounding mesenchyme at the limb sites, where they propagate and become committed to skeletal muscle progenitors (Relaix et al., 2005). By stimulating organoid culture at the neural plate border/dermomyotomal stage with bFGF/HGF we could further visualize both migration of myogenic progenitors and migration/specification of neural crest populations (**Figures 1A,G, H, 2A).** Further, by omitting FGF during organoid development, we could detect a continuous upregulation of genes involved in the myogenic migration process, such as LBX1, PAX3, PAX7 and CXCR4, but not in genes characterizing neural tube or neural crest development, such as SOX10, TFAP2A, PAK3, DLX5, B3GAT1, FOXB1, HES5 and OLIG3. This indicates that organoid culture conditions and specifically stimulation with HGF favored skeletal muscle over neural lineage **(Figures 2C-E)**. Interestingly, we could show that by stimulating organoid culture with SF/HGF, an essential signal for detachment and subsequent migration, but not for specification of the cells at the lateral lip of the dermomyotome (Dietrich et al., 1999), we could preserve the PAX3+/PAX7+ myogenic migratory population in an undifferentiated and uncommitted state **(Figure 2D,E).** Strikingly, expression profiling based on HOX gene cluster supported this notion, as over time the organoid culture adopted more distal than proximal limb axial anatomical identify **(Figure 2F)**.

Fetal development is characterized by PAX3+ to PAX7+ myogenic progenitor transition (Seale et al., 2000; Relaix et al., 2005), which we were able to demonstrate in our organoid culture. Our data further support organoid culture representing fetal stages as we could detect NFIX upregulation and verify presence of myofibers expressing fetal MyHC isoform as well as NCAM, M-Cad or MCAM in the proximity of PAX7 progenitors **(Figure 3G to J)**. Consequently, transcriptome comparison indicated high similarity to human fetal muscle tissue (17-week, Pearson correlation, rho=0.9) **(Figure 4E)**, as well as expression of several satellite cell markers **(4-figure supplement 1A).** This was further verified by comparison to a single cell dataset from the human skeletal muscle cell atlas **(Figures 4, 4-figure supplement 4)** Interestingly, single cell resolution showed adult satellite cell clusters with organoid derived myogenic progenitors **(Figure 4F**). In addition, pseudotime ordering indicated that organoid derived myogenic progenitors reside developmentally upstream of activated satellite cells, upregulating markers associated with quiescence such as NOTCH signaling and extracellular matrix proteins **(4-figure supplement 2E,F)**. Preservation of myogenic progenitors in a non-dividing state without activating MYOD1 and self-renewal **(Figures 4D, 4-figure supplement 2E,F))** appeared to be responsible for the observed sustainable propagation of PAX7 progenitor cells even 14 weeks post differentiation **(Figures 2F, 3A).**

Patterning in our 3D protocol provide progenitors in a more mature late fetal state partially overlapping with adult satellite cell developmental scoring **(Figure 4H)**. The ‘uncommitted’ Pax7 progenitor status is demonstrated by our precursor populations bulkRNAseq and scRNAseq profiling **(Figure 4 A,E, F)**. In this context, we could observe high expression of extracellular matrix proteins and upregulated NOTCH signaling in dormant non-dividing myogenic progenitors **(4-figure supplement 2A,E)**. This phenotype is similarly described for human fetal tissue myogenic progenitors (**Figure 4E**, Pearson correlation, rho=0.97**, 4-figure supplement 5)**. Studies evaluating engraftment potential of satellite cells and fetal muscle progenitors propose that muscle stem cells in a quiescent non-committed state exhibit enhanced engraftment potential (Hicks et al., 2017; Quarta et al., 2016; Montarras et al., 2005; Tierney et al., 2016). Our data demonstrate that upon activation and commitment dormant myogenic progenitors downregulate extracellular matrix proteins and upregulate expression of morphogens / receptors that make them susceptible to signals, like VEGFA that communicates with vasculature during tissue reconstitution, or CD9, CD44, CD98 participating in cellular activation (Porpiglia et al., 2017) **(2-figure supplement 1B,D)**. Cell-cell communication analysis revealed that myogenic progenitors influence their own fate mainly with ECM related signals (**4-figure supplement 5**). This finding is in line with the nature of in vivo fetal myogenic progenitors (Tierney et al., 2016) but also indicates that investigation at the myogenic progenitor level could provide new insights into ECM related congenital muscular dystrophies. One example is the Ullrich congenital muscular dystrophy, where the ECM alters the muscle environment with progressive dystrophic changes, fibrosis and evidence for increased apoptosis (Bönnemann, 2011).

CD82+ populations from our organoids engraft in the *Tibialis anterior* muscle of immunodeficient mice (**Figure 4I,J**). It would be of interest for future studies to investigate whether increased engraftment can be achieved in 3D protocols (Faustino Martins et al., 2020; Shahriyari et al., 2022; ours) versus 2D patterned progenitor cells and to which degree this is attributed to high expression of extracellular matrix proteins. In particular, high Fibrillin1 expression on dormant non-dividing myogenic progenitors could potentially contribute to avoidance of fibrosis by myogenic progenitors through regulation of TGF-β signaling (Cohn et al., 2007).

Different phases of human fetal myogenesis have been modelled with 2D differentiation protocols (Shelton et al., 2014, Chal et al. 2015, Xi et al. 2017). Our 3D differentiation protocol does not go beyond these protocols when it comes to provide maturated physiologically responsive skeletal muscle cells, which we illustrate with the electrophysiological recording of organoid-derived cells of different origins (**Figures 3 K,L,M, 3-figure supplement 3E,F**). Structural distinctions like the posterior paraxial mesoderm at Day 5, specified neural crest dermomytome at Day 17, myogenic progenitor migration at Day 23 and neural crest lineage arrest at Day 35 (**Figures 1 C-H, 2 C**) cannot be similarily observed in 2D protocols. In addition, our 3D organoid protocol provides myogenic progenitors in dormant and activated states for at least 14 weeks in culture. We demonstrate that organoid culture sustains uncommitted MyoD1-negative, Pax7-positive myogenic progenitors as well as fibroadipogenic (PDGFRa+) progenitors, both resembling their fetal counterpart. This supply of muscle progenitors preserved in a quiescent state indicates translative potential of the approach. Future work will elucidate signaling transduction pathways during skeletal muscle organoid development to model and understand human myogenesis in more detail.

## Materials and Methods

### hiPSCs culture

Human induced pluripotent stem cell (hiPSC) lines, Cord Blood iPSC (CB CD34^+^, passage 15 – 35), Duchenne Muscle Dystrophy iPSC patient line iPSCORE_65_1 (WiCell, cat.no. WB60393, passage 22-30) and DMD_iPS1 (passage 21-30) & BMD_iPS1 (passage 17-25) (Boston Children’s Hospital Stem Cell Core Facility), LGMD2A iPSC and LGMD2A-isogenic iPSC (Dorn et al., 2015; Mavrommatis et al., 2020; Panopoulos et al., 2017; Park et al., 2008) were cultured in TESR-E8 (StemCell Technologies) on Matrigel GFR (Corning) coated 6 well plates. The study was performed after ethical approval from the ethics commission of the Ruhr-University Bochum, Medical Faculty (15-5401, 08/2015).

### Human skeletal muscle organoid differentiation protocol

Prior differentiation, undifferentiated human PSCs, 60-70% confluent, were enzymatically detached and dissociated into single cells using TrypLE Select (ThermoFisher Scientific). Embryoid bodies formed via hanging drop approach with each droplet containing 3-4×10^4^ human single PSCs in 20 μl were cultured hanging on TESR-E8 supplemented with Polyvinyl Alcohol (PVA) at 4mg/ml (SigmaAldrich) and rock inhibitor (Y-27632) at 10μM (StemCell Technologies) at the lid of Petri dishes. At the beginning of skeletal muscle organoid differentiation, embryoid bodies at the size of 250-300μm were embedded into Matrigel and cultured in DMEM/F12 basal media (ThermoFisher Scientific) supplemented with Glutamine (ThermoFisher Scientific), Non Essential Amino Acids (ThermoFisher Scientific), 100x ITS-G (ThermoFisher Scientific), (Basal Media) 3μM CHIR99021 (SigmaAldrich) and 0.5μM LDN193189 (SigmaAldrich). At Day 3, human recombinant basic Fibroblast Growth Factor (bFGF) (Peprotech) at 10ng/μl final concentration was added to the media. Subsequently, at Day 5 the concentration of bFGF was reduced to 5ng/μl and the media was further supplemented with 10nM Retinoic Acid (SigmaAldrich). The differentiation media, at Day 7, was supplemented only with human recombinant Sonic hedgehog (hShh) (Peprotech) at 34ng/μl, human recombinant WNT1A (Peprotech) at 20ng/μl and 0.5μM LDN193189. At Day 11 the cytokine composition of the media was changed to 10ng/μl of bFGF and human recombinant Hepatocyte Growth Factor (HGF) at 10ng/μl (Peprotech). From Day 15 onwards, the basal media was supplemented with ITS-X (ThermoFisher Scientific) and human recombinant HGF at 10ng/μl. The first 3 days of the differentiation the media was changed on a daily basis, from 3^rd^ till 30^th^ every second day, while from Day 30 onwards every third day. The organoid approach was evaluated with six hiPSCs lines with independent genetic backgrounds, with more than five independent derivations per line, especially for the control line (CB CD34^+^) more than 20 derivation, obtaining always similar results. Per derivation and upon embryoid body Matrigel embedding, cultures exhibited high reproducibility. Upon migration and skeletal muscle formation organoids are occupying whole Matrigel droplet and through generation of additional bulge reach sizes of 4-5 mM. Concomitantly, myogenic progenitors fall off the organoid and generate a sheath of myogenic organoids and muscle fibers at the surface of the culture plate. For all lines, functional myofibers and PAX7 positive myogenic populations could be evaluated. Myogenic populations from different lines exhibit high similarity, (Pearson correlation, rho = 0.94-0,95, Figure 4 - figure supplement 3G).

### Immunocytochemistry

#### Cryosection Immunochemistry

Organoids from different stages were fixed on 4% paraformaldehyde overnight at 4°C under shakings conditions, dehydrated (30% sucrose o/n incubation) and embedded in OCT freezing media. Cryosections were acquired on a Leica CM3050s Cryostat. For the immunostaining process, cryosections were rehydrated with PBS and followed by permeabilization once with 0.1% Tween-20 in PBS, (rinsed 3x with PBS), and then with 0.1% Triton-X in PBS (rinsed 3x with PBS). Subsequently, the sections were blocked with 1% BSA/ 10% NGS in PBS for 1hr at room temperature. Primary antibody incubations were performed from 1 to 2 days at 4°C, where secondary antibody incubations for 2hr at room temperature.

#### EdU staining

At 12-week post differentiation, control organoids were incubated with 2.5μM BrdU final concentration overnight. To detect EdU, the sections were processed with Click-iT EdU Alexa Fluor 488 cell proliferation kit (Invitrogen) following the manufacturer’s instructions. The samples were incubated with secondary antibodies after the click reaction for detecting EdU. *Primary Antibodies*: anti-Brachyury (R&DSystems, 1:250), anti-TBX6(Abcam, 1:200), anti-PAX3 (DHSB, 1:250), anti-PAX7(DHSB, 1:250), anti-SOX10 (R&DSystems,1:125), anti-KI67 (ThermoFisher Scientific, clone SolA15, 1:100), anti-TITIN (DHSB, 9D-10, 1:300), anti-MyHC (DHSB, MF20, 1:300), anti-MYOD1 (Santa Cruz Biotechnologies, clone 5.8A, 1:200), anti-PRDM16 (Abcam, ab106410, 1:200), anti-TFAP2A (DHSB, 3B5, 1:100), anti-Dystrophin (Novocastra/Leica Biosystems, clone DY4/6D3, 1:200), anti-Laminin (SigmaAldrich, 1:200), anti-FastMyHC (SigmaAldrich, clone MY-32, 1:300), anti-M-Cadherin (Cell Signaling Technology, 1:200) anti-SOX2 (ThermoFisher Scientific, clone Btjce, 1:100), anti-CD44 (eBioscience, clone IM7, 1:100), anti-Fibrillin1 (Invitrogen, clone 11C1.3, 1:100).

#### Secondary antibodies

Alexa Fluor® 647 AffiniPure Fab Fragment Goat Anti-Mouse IgM, μ Chain Specific (Jackson Immunoresearch Laboratories, 1:100), Rhodamine RedTM-X (RRX) AffiniPure Goat Anti-Mouse IgG, Fcγ Subclass 1 Specific (Jackson Immunoresearch Laboratories,1:100), Alexa Fluor® 488 AffiniPure Goat Anti-Mouse IgG, Fcγ subclass 2a specific (Jackson Immunoresearch Laboratories,1:100), Alexa Fluor 488, Goat anti-Rat IgG (H+L) Cross-Adsorbed Secondary Antibody, (ThermoFisher Scientific, 1:500), Alexa Fluor 488, Donkey anti-Mouse IgG (H+L) Cross-Adsorbed Secondary Antibody, (ThermoFisher Scientific, 1:500), Alexa Fluor 647, Donkey anti-Goat IgG (H+L) Cross-Adsorbed Secondary Antibody, (ThermoFisher Scientific, 1:500), Alexa Fluor 488, Donkey anti-Goat IgG (H+L) Cross-Adsorbed Secondary Antibody, (ThermoFisher Scientific, 1:500), Alexa Fluor 568, Donkey anti-Rabbit IgG (H+L) Cross-Adsorbed Secondary Antibody, (ThermoFisher Scientific, 1:500). Images were acquired on a ZEISS LSM780 inverted confocal microscope.

### Oil O Red Staining

For histological visualization of adipocytes within the organoids, Oil O Red Stain kit (Abcam, ab150678) was applied on frozen sections derived from PFA fixated organoids following manufactures recommended protocol. Organoid sections upon staining with Oil O Red were visualized at an Olympus BX61 upright microscope.

### Flow Cytometry

#### FACS intracellular staining

Organoids during the 2^nd^,5^th^,9^th^ and 14^th^ week of differentiation were dissociated into single cells by incubating them till dissociating at 37°C within papain solution under shaking conditions. Then, the cells were pelleted at 400xg for 5min, followed by incubation with TryplE Select for 10min to ensure dissociation into single cells. Further, the cells were passed through 70μM (2^nd^ week) – 100μM (5^th^, 9^th^, 14t^h^ week) cell strainers to avoid aggregates. For both digesting steps 10% FBS/DMEM-F12 as digesting deactivation solution was applied to the cells. Then, for intracellular flow cytometric staining analysis the Transcription Factor Buffer set (BD Pharmigen) was applied and the cells were analyzed using flow cytometer (BD Biosciences FACS ARIAII). Primary antibodies used in this study: anti-PAX7, anti-MYOD1, anti-Pax3 in total amount of 400μg per staining; Secondary antibodies: Rhodamine RedTM-X (RRX) AffiniPure Goat Anti-Mouse IgG, Fcγ Subclass 1 Specific (Jackson Immunoresearch Laboratories), Alexa Fluor® 488 AffiniPure Goat Anti-Mouse IgG, Fcγ subclass 2a specific (Jackson Immunoresearch Laboratories) in 1:50 dilution. As isotype controls, Mouse IgG1 kappa Isotype Control (Invitrogen, clone P3.6.2.8.1), Mouse IgG2a kappa Isotype Control, (Invitrogen, clone eBM2a) were used at 400μg total amount per staining.

#### FACS EdU assay

At 15-week post differentiation, organoids were incubated overnight with 5μM EdU final concentration. Next day, organoids were dissociated into single cells by incubation at 37°C within papain solution for 1-2hr, followed by incubation with TryplE Select for 10min to ensure single cell dissociation. Then, the dissociated cells were passed through a 70μm cell culture strainer to remove any remaining aggregates. To detect EdU, the cells were processed with Click-iT EdU Alexa Fluor 488 Flow Cytometry Assay Kit (Invitrogen) according to manufacturer instructions and then analyzed using the 488 channel of a BD Biosciences FACSAria Fusion flow cytometer.

#### FACS isolation of ITGβ1^+^ / CXCR4^+^ myogenic cell population for RNA sequencing

Organoids from Duchenne and Control iPSC lines and during 15th - 16th week post differentiation were dissociated into single cells during incubation with Papain solution dissociation upon gentle shaking (1-2hr) was observed. To acquire singlets, the cells were filtered through 40μm cell strainers and upon washing with 1% BSA solution prepared for surface antigen staining. For surface antigen staining 20 min incubation with the Alexa Fluor 488 anti-human CD29 (Biolegend, clone TS2/16), PE anti-human CD184[CXCR4] (Biolegend, clone 12G5) was applied together with the corresponding isotype controls: PE Mouse IgG2a, κ Isotype Ctrl antibody (Biolegend, clone MOPC-173), 488 Mouse IgG1, κ Isotype Ctrl antibody (Invitrogen, clone P3.6.2.8.1). For removing residual antibodies, the cells were washed twice with 1% BSA staining solution and processed by BD Biosciences FACSAria Fusion flow cytometer. Briefly before FACS sorting to discriminate between dead and alive cells DAPI was added to the samples and then DAPI^-^ / CD29^+^ / CXCR4^+^ cells populations were collected into tubes containing RLT buffer supplemented with b-mercaptoethanol to avoid RNA degradation. The FACS gating strategy is further depicted in Figure 5 – figure supplement 3A.

### Bulk RNA sequencing

#### RNA extraction

Total RNA was extracted from single organoids or cultured cells by using the RNAeasy Micro Kit (Qiagen) according to the manufacturer’s instructions. Subsequently, before library preparation, the RNA integrity was evaluated on an Agilent 2100 Bioanalyzer by using the RNA 6000 Pico kit (Agilent). *cDNA library preparation*: For 4-week and 8-week organoids cDNA library was prepared by using the whole transcriptome Illumina TruSeq Stranded Total RNA Library Prep Kit Gold (Illumina), followed by evaluation on an Agilent 2100 Bioanalyzer by using the DNA 1000 kit. The resulting mRNA library was sequenced as 2 x 75 bp paired-end reads on a NextSeq 500 sequencer (Illumina). For 16-weeks and ITGβ1^+^ / CXCR4^+^ sorted cells cDNA library was prepared using the whole transcriptome Ovation Solo RNA seq Library Preparation Kit (TECAN, NuGEN), followed by evaluation on an Agilent 2100 Bioanalyzer by using the DNA 1000 Chip. The resulting mRNA library was sequenced as 1 x 150 bp single reads on a HiSeq. 3000 sequencer (Illumina).

### Bulk RNA seq bioinformatic analysis

Sequenced reads were aligned to the human reference genome (hg38) with TopHat2 (version 2.1.1), and the aligned reads were used to quantify mRNA expression by using HTSeq-count (version 0.11.2). DESeq2 (Love at al., 2014) was used to identify differentially expressed genes across the samples. ITGβ1^+^ / CXCR4^+^ organoid derived myogenic cell populations were compared to already available transcriptomic dataset of human fetal muscle progenitors (GSM2328841-2) (Hicks et al., 2018).

### scRNA sequencing

#### Sample and cDNA library preparation

Single cells were acquired upon incubation for 1hr with solution containing papain and EDTA. Upon dissociation the cell number and viability was estimated. Then cells were resuspended in a solution containing 0.5% BSA in PBS to reach a concentration of 390 cells per μl. The cDNA library was prepared using the Chromium Single Cell 3′ Reagent Kits (v3): Single Cell 3′ Library & Gel Bead Kit v3 (PN-1000075), Single Cell B Chip Kit (PN-1000073) and i7 Multiplex Kit (PN-120262) (10x Genomics) according to the manufacturer’s instructions. Then, the cDNA library was run on an Illumina HiSeq 3000 as 150-bp paired-end reads.

### Single cell RNA seq bioinformatic analysis

Sequencing data were processed with UMI-tools (version 1.0.0), aligned to the human reference genome (hg38) with STAR (version 2.7.1a), and quantified with Subread featureCounts (version 1.6.4). Data normalization and further analysis were performed using Seurat (version 3.1.3, Stuart et al., 2019). For initial quality control of the extracted gene-cell matrices, cells were filtered with parameters low threshold = 500, high threshold =6,000 for number of genes per cell (nFeature_RNA), high threshold = 5 for percentage of mitochondrial genes (percent.mito) and genes with parameter min.cells = 3. Filtered matrices were normalized by the LogNormalize method with scale factor = 10,000. Variable genes were found with parameters of selection.method = “vst”, nfeatures = 2000, trimmed for the genes related to cell cycle (KEGG cell cycle, hsa04110) and then used for principal component analysis. Statistically significant principal components were determined by the JackStraw method and the first 5 principle components were used for non-linear dimensional reduction (tSNE and UMAP) and clustering analysis with resolution=0.2. Monocle3 (version 0.2.0, Cao et al., 2019) was used for pseudotime trajectory analysis. The data matrix of Seurat objects (assays[[“RNA”]]@counts) was imported to the Monocle R package, then dimensionality reduction with the PCA method with parameters max_components=2 was performed and then cluster_cells, learn_graph and order_cells functions were performed subsequently. Organoid-derived myogenic progenitors were compared to already available transcriptomic dataset of adult satellite cells (GSE130646) (Rubenstein et al., 2020).

For integrative analysis we used the Seurat (version 4, Hao et al., 2021) package. For each dataset, cells with more than 6,000 or less than 300 detected genes, as well as those with mitochondrial transcripts proportion higher than 5-10% were excluded. For finding anchors between control and DMD datasets using 5000 anchors and regressing out cell cycle genes, sequencing depth and stress related genes was carried out before integration.

Developmental score was calculated as described in Xi et al. 2020. Briefly, we used the ‘‘AddModuleScore’’ function to calculate the embyonic and adult score using a list of differentially expressed genes (DEGs) between adult and embryonic myogenic progenitor clusters. DEGs were selected from the Supplementary information in Xi et al. 2020 table mmc3. The developmental score was further calculated by subtracting embryonic from the adult score. Embryonic and fetal datasets were filtered using a low threshold = 500 and adult datasets were filtered using a low threshold = 250 for genes per cell. In addition, scaling the data (‘‘S.Score,’’ ‘‘G2M.Score,’’ ‘‘Stress’’ and ‘‘total Count’’ to the vars.to.regress’ argument) was used to regress out the effects of the cell cycle, dissociation-related stress as well as cell size/sequencing depth to all datasets. Myogenic subpopulation of SMO and adult satellite cell datasets were selected by ‘PAX7’ expression. Embryonic and fetal (weeks 5 to 18) and adult satellite cell (years 7,11,34,42) scRNAseq data are from Xi et al., 2020 (GSE147457), adult satellite cell (year 25) scRNAseq data are from Rubenstein et al., 2020 (GSE130646).

### Cell-Cell communication analysis

To investigate cell-cell communications among activated, mitotic and dormant myogenic progenitors, fibroadipogenic and neural progenitors and myofiber related clusters from 12-week organoids the CellChat R package (Jin, S. et al., 2021) was applied.

### qPCR expression analysis

By pooling 3 organoids per sample, total RNA was extracted using the RNAeasy Mini Plus Kit (Qiagen). For first strand cDNA synthesis, the High capacity RNA to cDNA Kit (Applied Biosystems) was applied using as template 2μg of total RNA. For setting qPCR reactions, the GoTaq qPCR Master Mix (Promega) was used with input template 4ng cDNA per reaction while the reaction was detected on a CFX 96 Real-Time PCR detection system (BIO-RAD). The relative quantification (ΔCT) method was applied for detecting changes in gene expression of pluripotent, neural tube, neural crest, posterior/anterior somitic; and dermomyotomal markers, between different time points along the differentiation. qPCR primers applied for each marker for evaluating organoid development are listed in **Table S1**.

### Diffusion map analysis

By pooling 3-5 organoids per sample, total RNA was extracted using the RNAeasy Mini Plus Kit (Qiagen) and further processed as described under qPCR expression analysis. Normalized Ct values of selected genes for each sample was used as input for generating eigenvector values using the destiny package (Angerer, P. et al., 2016). Then, all samples were ordered by their Diffusion Component 1 at specific time-points during early culture development (Day 2 – Day 11). Ct values above 30 were not considered for subsequent analysis. qPCR primers applied for each marker for diffusion map analysis are listed in **Table S2**.

### Transmission Electron Microscopy (TEM)

Skeletal muscle organoids were fixed for 4h at room temperature (RT) in 2.5% glutaraldehyde (SigmaAldrich) in 0.1M cacodylate buffer pH 7,4 (Sciences Services, Germany), subsequently washed in 0.1M cacodylate buffer pH 7,4, post-fixed for 2h at RT in 1% osmium tetroxide (Sciences Services, Germany) in 0.1M cacodylate buffer pH 7,4, dehydrated stepwise in a graded ethanol series and embedded in Epon 812 (Fluka, Buchs, Switzerland). Ultrathin sections (70 nm, ultramicrotome EM UC7, Leica, Wetzlar, Germany) were afterwards stained for 30 min in 1% aqueous uranyl acetate (Leica, Germany) and 20 min in 3% lead citrate (Leica, Germany). TEM images were acquired with a 200 kV TEM JEM 2100Plus (Jeol, Japan), transmission electron microscope.

### Second harmonic generation (SHG) imaging using multi-photon microscopy

A TriM Scope II multi photon system from LaVision BioTec was used to visualize skeletal muscle fiber organization inside organoids and distinct sarcomeres. The microscope setup is a single beam instrument with an upright Olympus BX51 WI microscope stand equipped with highly sensitive non-descanned detectors close to the objective lens. The TriM Scope II is fitted with a Coherent Scientific Chameleon Ultra II Ti:Sapphire laser (tuning range 680-1080 nm) and a Coherent Chameleon Compact OPO (automated wavelength extension from 1000 nm to 1600 nm). A 20x IR objective lens (Olympus XLUMPlanFl 20x/1.0W) with a working distance of 2.0 mm was used. Muscle fiber SHG signals were detected in forward direction using TiSa light at 850 nm, a 420/40 band pass filter and a blue-sensitive photomultiplier (Hamamatsu H67080-01). 3D-images were acquired and processed with LaVision BioTec ImSpector Software.

### Electrophysiology

#### Current measurement

Membrane currents were measured at ambient temperature (22-24°C) using standard whole-cell patch clamp software ISO2 (MFK, Niedernhausen, Germany). Cells were voltage-clamped at a holding potential of –90 mV, i.e. negative to EnAChR, resulting in inward Na^+^ currents. Every 10 s, voltage ramps (duration 500 ms) from -120 mV to +60 mV were applied to assess stability of the recording conditions and to generate I/V curves (membrane currents in response to depolarizing voltage ramps are shown as downward deflections). Signals were filtered (corner frequency, 1 KHz), digitally sampled at 1 KHz and stored on a computer equipped with the hardware/software package ISO2 for voltage control, data acquisition and data analysis. Rapid exposure to a solution containing acetylcholine was performed by means of a custom-made solenoid-operated flow system permitting a change of solution around an individual cell with a half time of about 100 ms. For measurements cells devoid of contact with neighboring cells were selected. Cells originated from organoids at week 8.

#### Fluorescence microscopy and imaging

To monitor changes in [Ca2^+^]i, skeletal muscle cells were transiently transfected with pcDNA3[Twitch-2B] (Addgene, 49531) (0.25 μg per 35 mm culture-dish). Skeletal muscle cells were transfected using either poly-ethyleneimine (PEI) or Lipofectamine (Invitrogen) according to the manufacturer’s instructions. Prior to experiments, cells were seeded on sterile, poly-L-lysine-coated glass cover slips and analyzed or 48 h after transfections. All experiments were performed using single cells at ambient temperature.

Fluorescence was recorded buckusing an inverted microscope (Zeiss Axiovert 200, Carl Zeiss AG, Göttingen, Germany) equipped with a Zeiss oil immersion objective (100x/1.4), a Polychrome V illumination source and a photodiode-based dual emission photometry system suitable for CFP/YFP-FRET (FEI Munich GmbH, Germany). For FRET measurements, single cells were excited at 435 nm wavelength with light pulses of variable duration (20 ms to 50 ms; frequency: 5 Hz) to minimize photo-bleaching. Corresponding emitted fluorescence from CFP (F480 or FCFP) or from YFP (F535 or FYFP) was acquired simultaneously and FRET was defined as ratio FYFP/FCFP. Fluorescent signals were recorded and digitized using a commercial hardware/software package (EPC10 amplifier with an integrated D/A board and Patch-master software, HEKA, HEKA Elektronik, Germany). The individual FRET traces were normalized to the initial ratio value before agonist application (FRET/FRET0).

#### Solutions and chemicals

For FRET measurements an extracellular solution of the following composition was used (mmol/L): NaCl 137; KCl 5.4; CaCl2 2; MgCl2 1.0; Hepes/NaOH 10.0, pH 7.4. For whole cell measurements of membrane currents an extracellular solution of the following composition was used (in mmol/L): NaCl 120, KCl 20, CaCl2 0.5, MgCl2 1.0, HEPES/NaOH 10.0 (pH 7.4). The pipette solution contained (in mmol/L): K-aspartate 100, KCl 40, NaCl 5.0, MgCl2 2.0, Na2ATP 5.0, BAPTA 5.0, GTP 0.025, and HEPES/KOH 20.0, pH 7.4. Standard chemicals were from Merck. EGTA, HEPES, Na2ATP, GTP and acetylcholine chloride, were from SigmaAldrich.

### Transplantation experiments

Flow cytometry: The organoid culture was dissociated using TrypLe (Thermo Fisher Scientific, 12563011) and were filtered through 70 - 40 μm cell strainer. Single cell suspension was stained with PE anti-human CD82 Antibody (BioLegend, 342103) in 2%BSA and 2mM EDTA and sorted using FACS sorter (Beckman Coulter, MoFlo Astrios Zellsortierer). For gating, unstained single cells from the same organoid culture were used as a baseline control. Sorted cells were further processed for transplantation experiments.

Transplantation of skeletal muscle progenitors into murine *Tibialis anterior* (TA) muscle: 12 week old organoids were dissociated and single cells were sorted using FACS (CD82+). Cell transplantation was carried out as described before (Alexander et al., 2016, Marg et al., 2019, Al Tanoury et al., 2020). Briefly, 24h before transplantation, Cardiotoxin (10 µl CTX, 40ng/ml, Sigma-Aldrich, 217503-1MG) was injected into TA of 2-3 months old male HsdCpb:NMRI-Foxn1nu mice. Under anesthesia 1*10^5^ cells were injected into the TA muscle on one side. 6 weeks after transplantation the mice were killed and the TA were fixed and sliced using cryosections. Immunofluorescence analyses were carried out using recombinant Anti-Lamin A + Lamin C antibody and anti-Dystrophin antibody.

All animal experiments were approved by the local authorities (81-02.04.2020.A476, Ruhr University Bochum) and performed in accordance with the guidelines for Ethical Conduct in the Care and Use of Animals.

Cryosection Immunochemistry: TA from mice were fixed on 4% paraformaldehyde overnight at 4°C and kept in increasing sucrose solutions (10%, 20%, 30%) at 4 °C until the tissue sinked down (usually 24h) and embedded in Tissue-Tek® O.C.T.™ Compound media. Cross-sections of TA with 10 - 20 μm thickness were acquired on CryoStar NX50 (Thermo Scientifc). Sections were stored on objectives at -20°C.

Cryosections were rehydrated with PBS and followed by an antigen retrieval, an AffiniPure Fab Fragment Goat Anti-Mouse IgG treatment to prevent unspecific bindings during the staining process and permeabilized once with 1% (vol/vol) Triton-X100 and 125 mM glycine in PBS, (rinsed 3x with PBS). Subsequently, the sections were blocked with 5% BSA / 10% NGS in PBS for 1hr at room temperature. Primary antibody incubations were performed for 1 day at 4°C and secondary antibody incubations for 2hr at room temperature.

Primary Antibodies: anti-Dystrophin (Leica, NCLDYS1, 1:20), recombinant Anti-Lamin A + Lamin C antibody (Abcam, ab108595, 1:150) Secondary antibodies: Goat anti-Rabbit IgG (H+L) Highly Cross-Adsorbed Secondary Antibody, Alexa Fluor Plus 488 (1:1000), Goat anti-Mouse IgG (H+L) Cross-Adsorbed Secondary Antibody, Alexa Fluor 568 (1:1000)

Images were acquired on a Zeiss Scan.Z1. Images were processed using Zen Lite Blue version 4.0.3..

### Statistics

All statistical analysis was conducted using GraphPad Prism6 software. For qPCR analysis one-way ANOVA with Tukey ′s multiple comparisons test for each marker was performed. For the FACS intracellular staining quantification, one-way ANOVA with Sidak ′s multiple comparisons test between the different time points was performed. Significance asterisks represent *P < 0.05, **P < 0.01, ***P < 0.001, ****P < 0.0001, ns: not significant.

### Data and code availability

RNA sequencing datasets produced in this study are deposited in the Gene Expression Omnibus (GEO) under accession code GSE147514. Detailed scripts and parameters used for the study are available from the authors upon reasonable request.

**To review GEO accession GSE147514: Go to** https://www.ncbi.nlm.nih.gov/geo/query/acc.cgi?acc=GSE147514

## ACKNOWLEDGEMENTS

We are grateful to Drs. Karl Köhrer, Tobias Lautwein, Patrick Petzsch and Thorsten Wachtmeister, Genomics & Transcriptomics Laboratory, Heinrich-Heine-University Düsseldorf for performing single cell and bulk RNAseq experiments with their Illumina HiSeq platform and data provision. We are further grateful to Dr. Oliver Griesbeck for the pcDNA3[Twitch-2B] plasmid. We would like to thank Ingrid Gelker and Martina Sinn, Max Planck Institute Münster as well as Eva-Maria Konieczny, Rana Houmany and Boris Burr, Ruhr-University Bochum for their technical assistance. We thank Dr. Johnny Kim, Max Planck Institute Bad Nauheim for discussions and Dr. Elisabeth Stevens, *English Scientific*, Düsseldorf for scientific editing of the manuscript. Electron microscopy experiments were supported by the Deutsche Forschungsgemeinschaft SFB 944. We thank Drs. George Q. Daley and Thorsten Schlaeger, Boston Children’s Hospital for providing the Duchenne Muscular Dystrophy patient-derived iPS cell lines DMD-iPS1 and DMD-iPS2 in the course of our study. Our study was supported by research grants from FoRUM F873-16, Medical Faculty, Ruhr University Bochum, from Deutsche Gesellschaft für Muskelkranke e.V. (DGM Foundation), Freiburg, Georg E. und Marianne Kosing-Stiftung, Deutsches Stiftungszentrum, Essen and Deutsche Duchenne Stiftung, Duchenne Deutschland e.V..

## AUTHOR CONTRIBUTIONS

L.M. conceived and designed study, designed and performed experiments, single cell bioinformatic analyses, analyzed data, and wrote manuscript, H.W.J., U.K., D.G., J.H.Y., M.J.A.B. performed (sc)RNAseq experiments and analyzes, U.K., G.G.G., M.C.K., M.S., O.E.P., D.Z., M.G.B., D.H., G.M.P. performed experiments and analyzed data, J.B.K., J.C.S., S.H.A, R.H.A., H.R.S. provided study materials and supervised experiments, M.V. provided study materials, biopsies and supervised experiments, B.B.S. provided study materials, supervised experiments and edited manuscript, H.Z. designed and supervised study, experimental design, provided study materials and wrote manuscript.

---

**Figure 1 – figure supplement 1.**
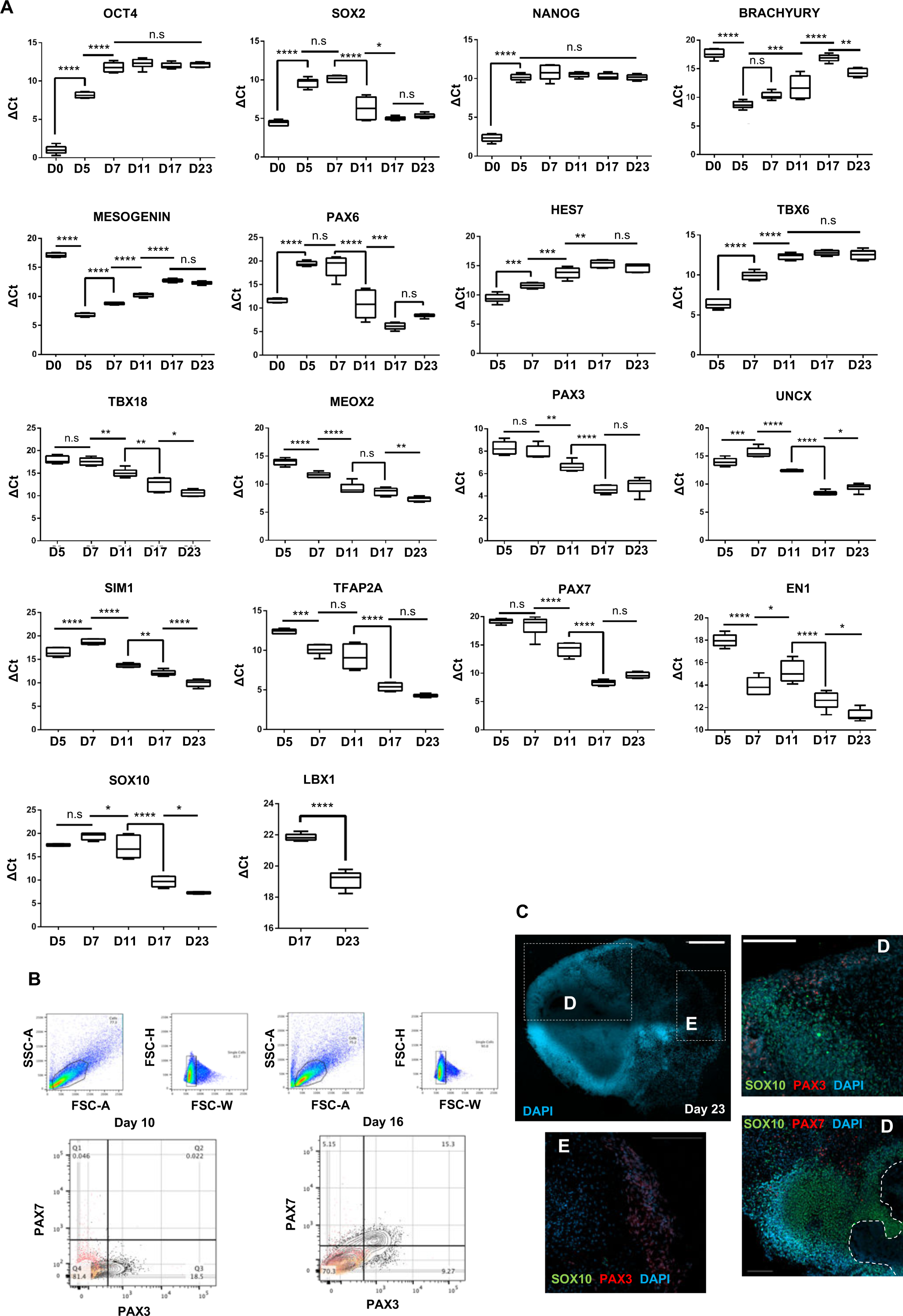
Lineage representation and organoid culture progression at early stages of differentiation protocol. **(A)** qPCR analysis depicts relative expression of pluripotent (OCT4, SOX2, NANOG), mesodermal / posterior somitic (BRACHYURY,MESOGENIN,TBX6,HES7), anterior somitic (PAX3,UNCX,TBX18,MEOX2), dermomyotomal (PAX7, SIM1, EN1, LBX1) and neural tube / crest markers (SOX2, PAX6, TFAP2A, SOX10) **(B)** Gating strategies and FACS intracellular quantification of PAX3 and PAX7 markers at Day 10 and 16 **(C)** Tile scan overview at Day 23 shows neural crest SOX10+ populations close to neural plate border epithelium **(D)** and SOX10-/PAX3+ populations at more outer locations **(E)**. For each replicate 6 organoids were pooled; gene expression was normalized to housekeeping gene. Statistics: *P < 0.05, **P < 0.01, ***P < 0.001, ****P < 0.0001, ns: not significant. FACS plots, yellow: unstained population, red: isotype control, gray: PAX7+ or PAX3+ population. Scale bars, 500uM (C), 200uM (D, E)

**Figure 2 – figure supplement 1.**
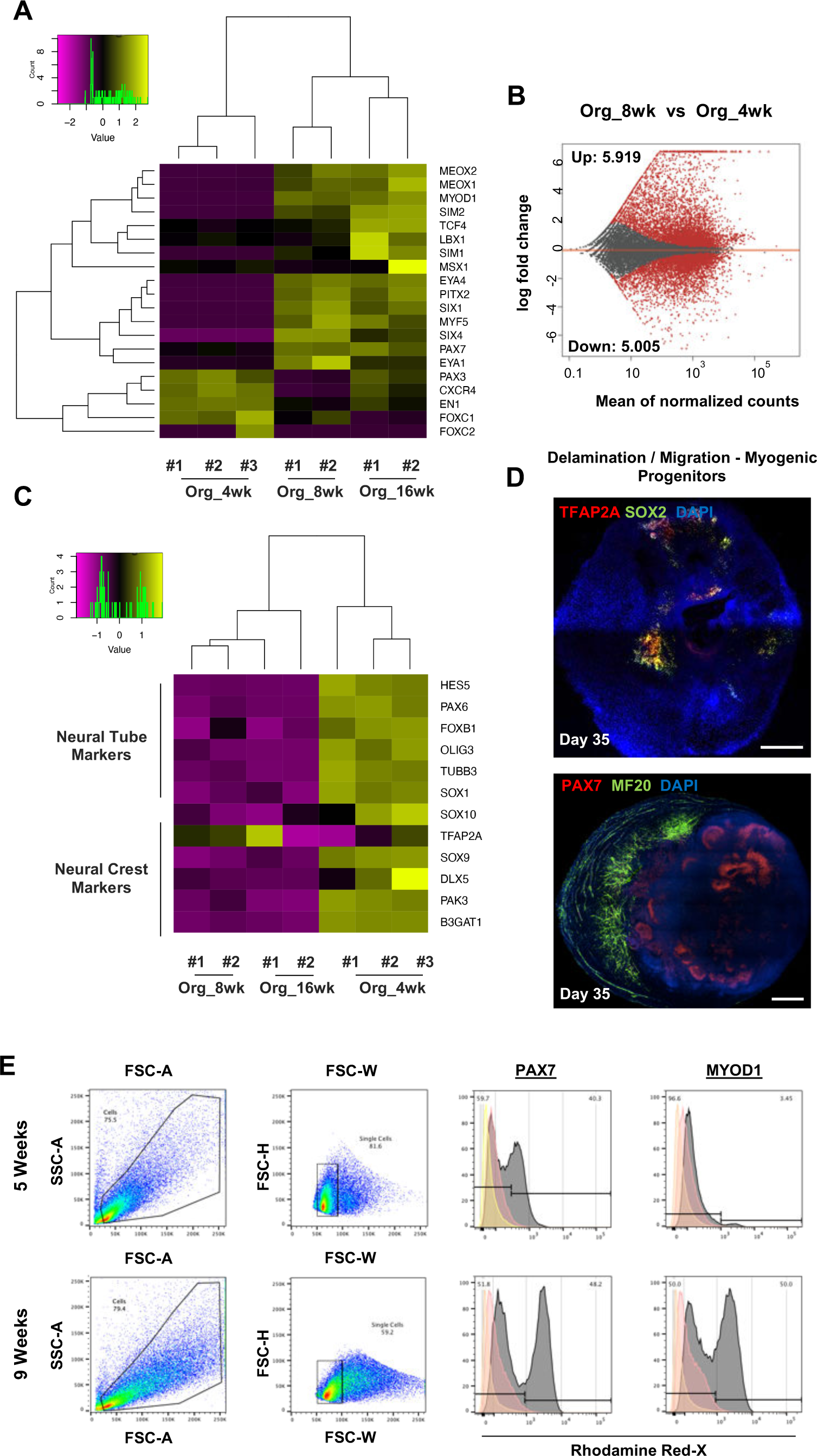
Myogenic versus neural fate during organoid development. **(A)** Heatmap of limb myogenic progenitor markers emphasizes upregulation through organoid protocol development (**B)** Differential expression comparison highlights upregulated (*n*=5.919) and downregulated (*n*=5.005) genes between 8 vs 4 weeks organoids. (Padj< 0.001) **(C)** Heatmap of neural tube and neural crest markers during 4, 8 and 16 weeks post differentiation highlights neural lineage arrest from 4 weeks onwards **(D)** Organoid overview at Day 35 indicates presence of MF20^+^ myofibers in proximity of PAX7^+^ cells and delamination migration of mainly myogenic cells. SOX2 and TFAP2A expression is restricted towards inner organoid areas **(E)** Gating strategy applied to quantify MYOD1^+^ (committed state) and PAX7^+^ (progenitor state) populations at 5 and 8 weeks post differentiation. Yellow histogram: unstained population, red: isotype control, gray: PAX7^+^ or MYOD^+^ population. Scale bars, 500uM

**Figure 3 – figure supplement 1.**
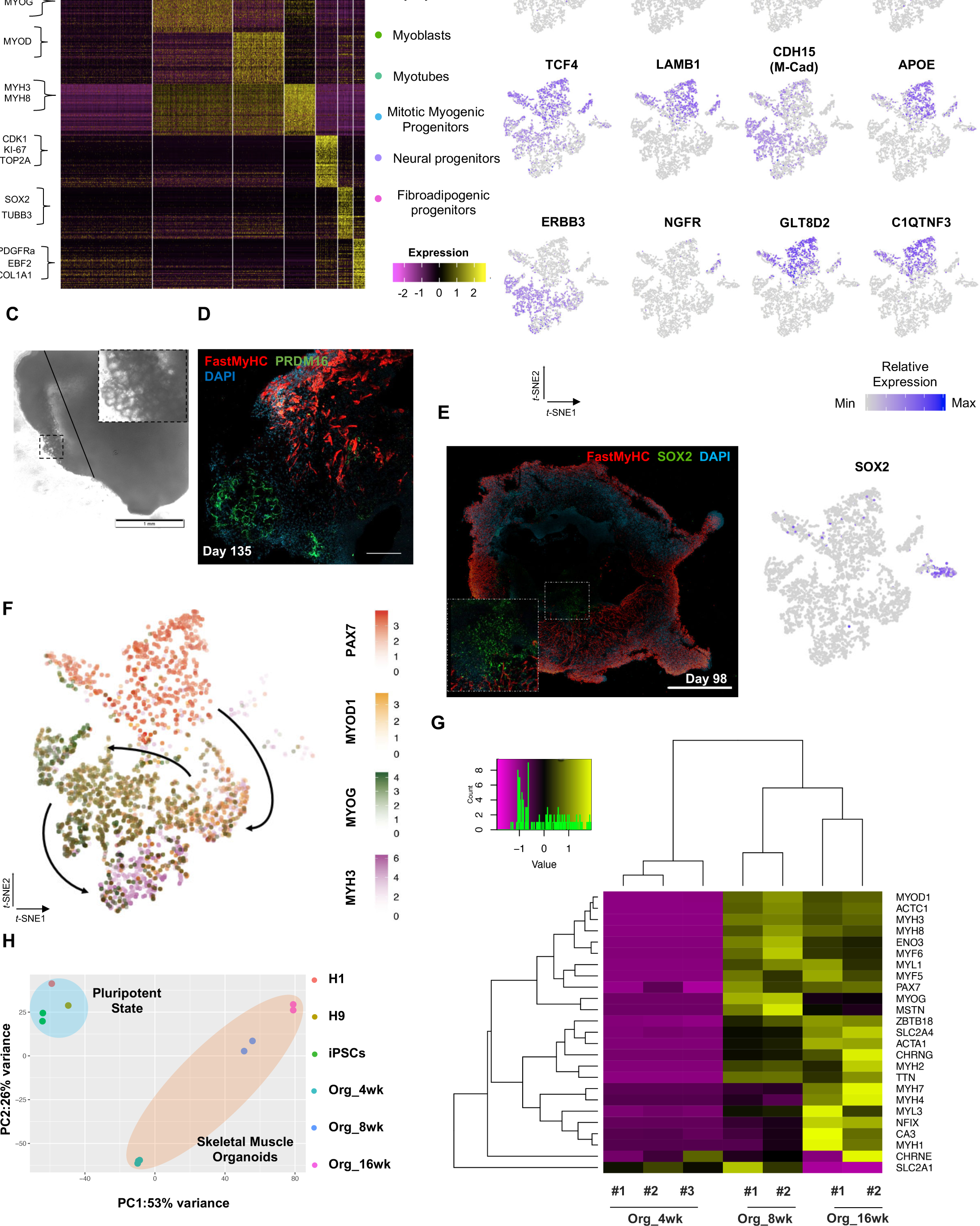
Single cell RNA seq expression profiling and lineage representation in organoid culture at week 12. **(A)** Gene signatures of *t*-SNE described clusters based on relative expression levels of the 50 most significant markers for each of the 7 clusters **(B)** *t*-SNE plot of myogenic and fibroadipogenic markers **(C)** Brightfield image at Day 135 highlights presence of structures reassembling adipocytes **(D)** Immunocytochemistry at Day 135 emphasizing derivation of PRDM16^+^ cells in FastMyHC^+^ myofibers proximity **(E)** Organoid overview at Day 98 highlights muscle system development at outer and neural lineage representation at more inner sites **(F)** *t*-SNE plot of key markers PAX7, MYOD1, MYOG, MYH3 from each stage depicts relative expression levels and emphasize gradual transition from myogenic progenitor to myotube subcluster **(G)** Heatmap of embryonic or fetal myogenic markers depicts a fetal environment during 8 to16 weeks post differentiation **(H)** Two-dimensional principal component analysis based on gene expression profiling between samples separates skeletal muscle organoid cluster from pluripotent and highlights greater difference in variance between 4 and 8-16 weeks subgroups in comparison to 8 and 16 weeks subgroups. w.p.d.: weeks post differentiation. Scale bars, 1mM (D), 200uM (E, F).

**Figure 3 – figure supplement 2.**
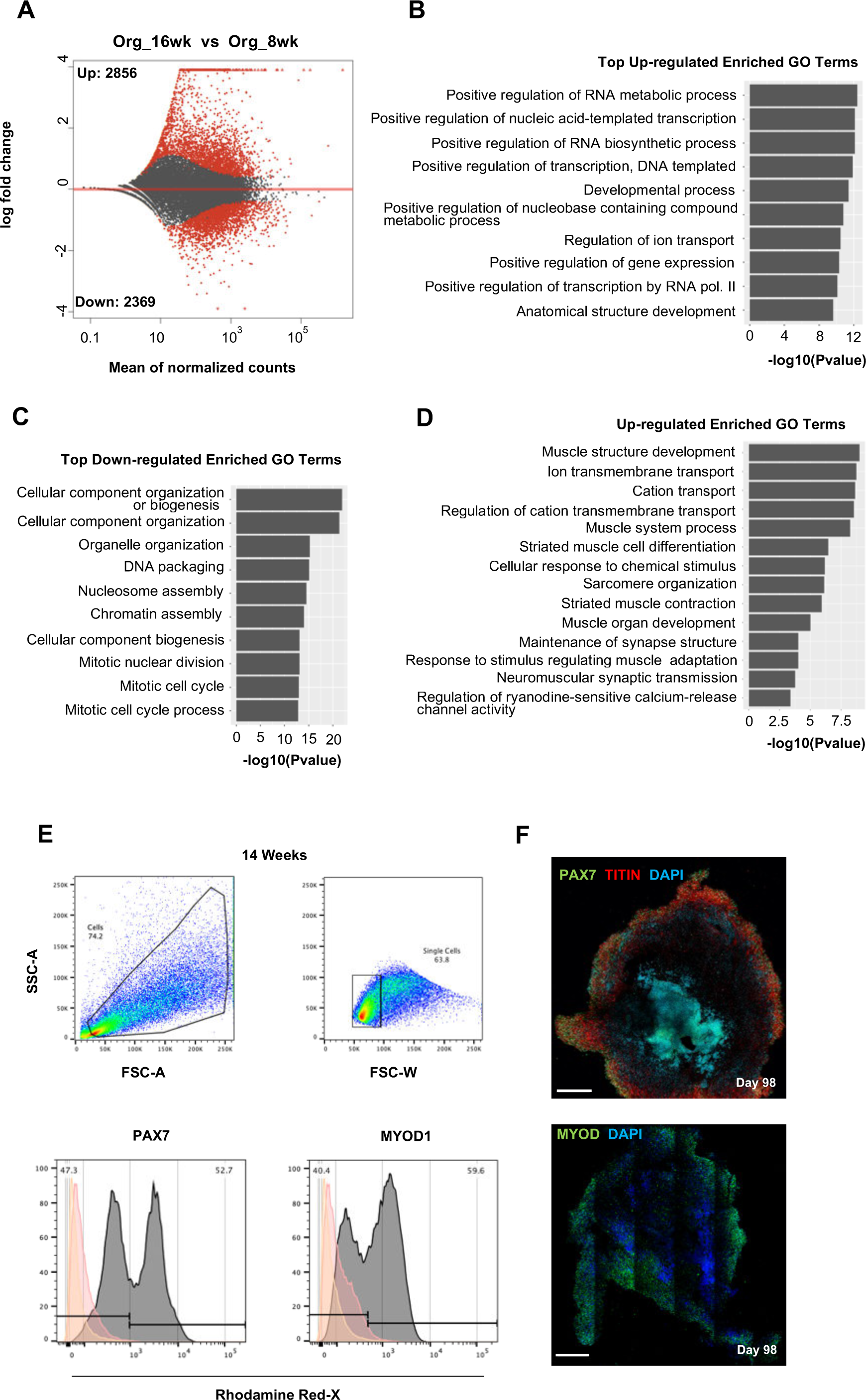
Skeletal muscle organoid culture maturation and identity. **(A)** Differential expression comparison highlights upregulated (*n*=2.856) and downregulated (*n*=2.369) genes between 16 and 8 weeks organoids (Padj<0.001) **(B-C)** Gene ontology enrichment analysis at 16 weeks depicting top statistically significant upregulated **(B)** and downregulated **(C)** gene ontology (GO) terms; **(D)** statistically significant upregulated GO terms further highlights skeletal muscle maturation **(E)** Gating strategy applied to quantify MYOD1^+^ (committed state) and PAX7^+^ (progenitor state) population at 14 w post differentiation **(F)** Organoid overview at Day 98 depicts high proportion of PAX7^+^ /MYOD^+^ cells towards outer areas. Yellow histogram: unstained population, Red histogram: Isotype Control, Gray Histogram: PAX7^+^ or MYOD^+^ population. Scale bars, 500uM (F)

**Figure 3 – figure supplement 3.**
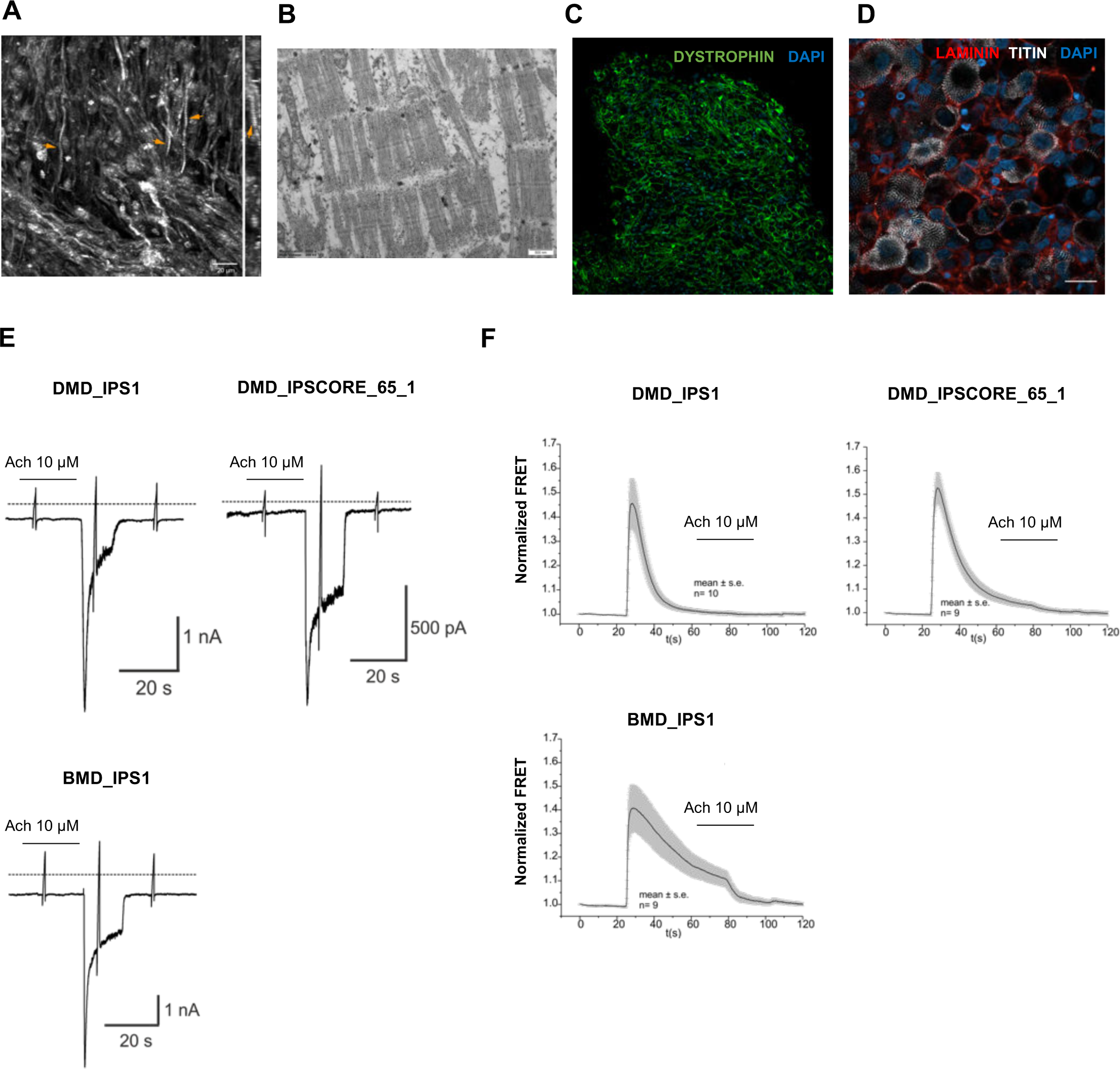
Functional properties of organoid derived skeletal muscle myofibers. **(A)** Second Harmonic Generation imaging (SHG) from areas with dense fiber network reveals myofibril formation with distinct sarcomeres (z plane of stack right) **(B)** Electron microscopy depicts myofibrils with well-developed sarcomeres **(C,D)** Skeletal muscle organoid derived myofibers expressed LAMININ and DYSTROPHIN **(E)** Representative recording of acetylcholine (Ach)-induced changes in holding current in a single skeletal muscle cell from organoid derived skeletal muscle cells from iPSC lines with Duchenne (DMD_IPS1, DMD_IPSCORE_65_1) and Becker (BMD_IPS1) muscular dystrophy patients. ACh (10 μM) was applied as indicated by the bar. Holding potential -90 mV. Downward deflections represent membrane currents in response to depolarizing voltage ramps (duration 500 ms) from -120 mV to +60 mV. Dashed line indicates zero current level. **(F)** Summarized FRET-recordings from organoid derived skeletal muscle cells from iPSC lines with Duchenne (DMD_IPS1, DMD_IPSCORE_65_1), Becker (BMD_IPS1) transfected with Twitch2B to monitor the increase in [Ca2+]i during ACh application. Scale Bars, 100uM (C,D), 20uM in (A), 500nM (B)

**Figure 4 – figure supplement 1.**
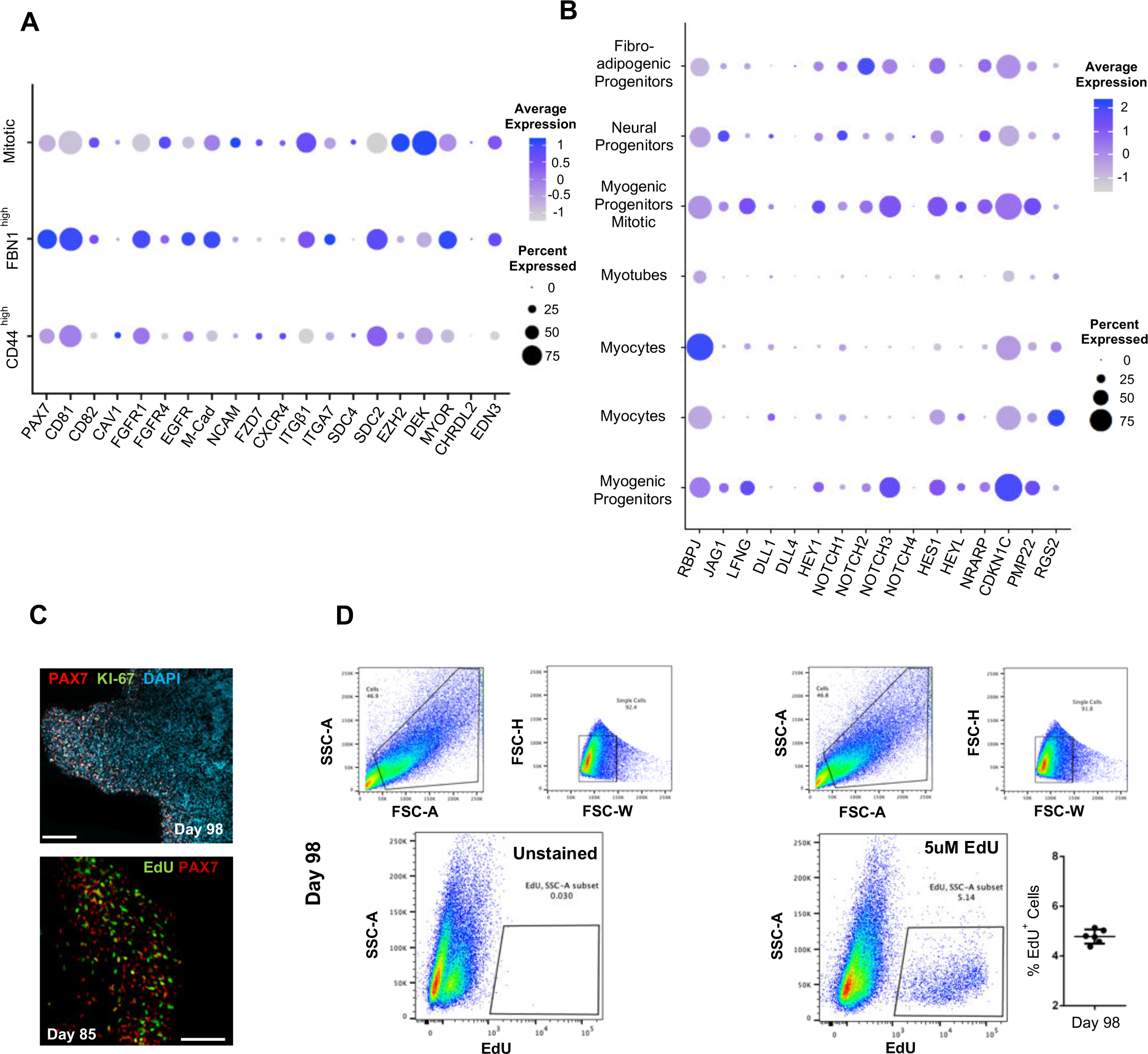
Subclustering of myogenic progenitors and NOTCH signaling. **(A)** Dot plot showing expression of representative genes related to satellite cell identity across the 7 main clusters. **(B)** Dot plot showing expression of representative genes related to NOTCH signaling, which are identified as regulators of satellite cell quiescence across the 7 main clusters **(C)** Organoid immunohistochemistry at Day 98 indicates PAX7 myogenic progenitors that are Ki-67-/EdU-**(D)** Gating strategy applied to quantify EdU+ proliferating cells in organoid culture at Day 98 (14 weeks); Histograph depicting percentage of EdU+ cells; Dot plot, circle area represents percentage of gene+ cells in a cluster, color reflects average expression level (gray, low expression; blue, high expression). Scale Bars, 100uM in (C).

**Figure 4 – figure supplement 2.**
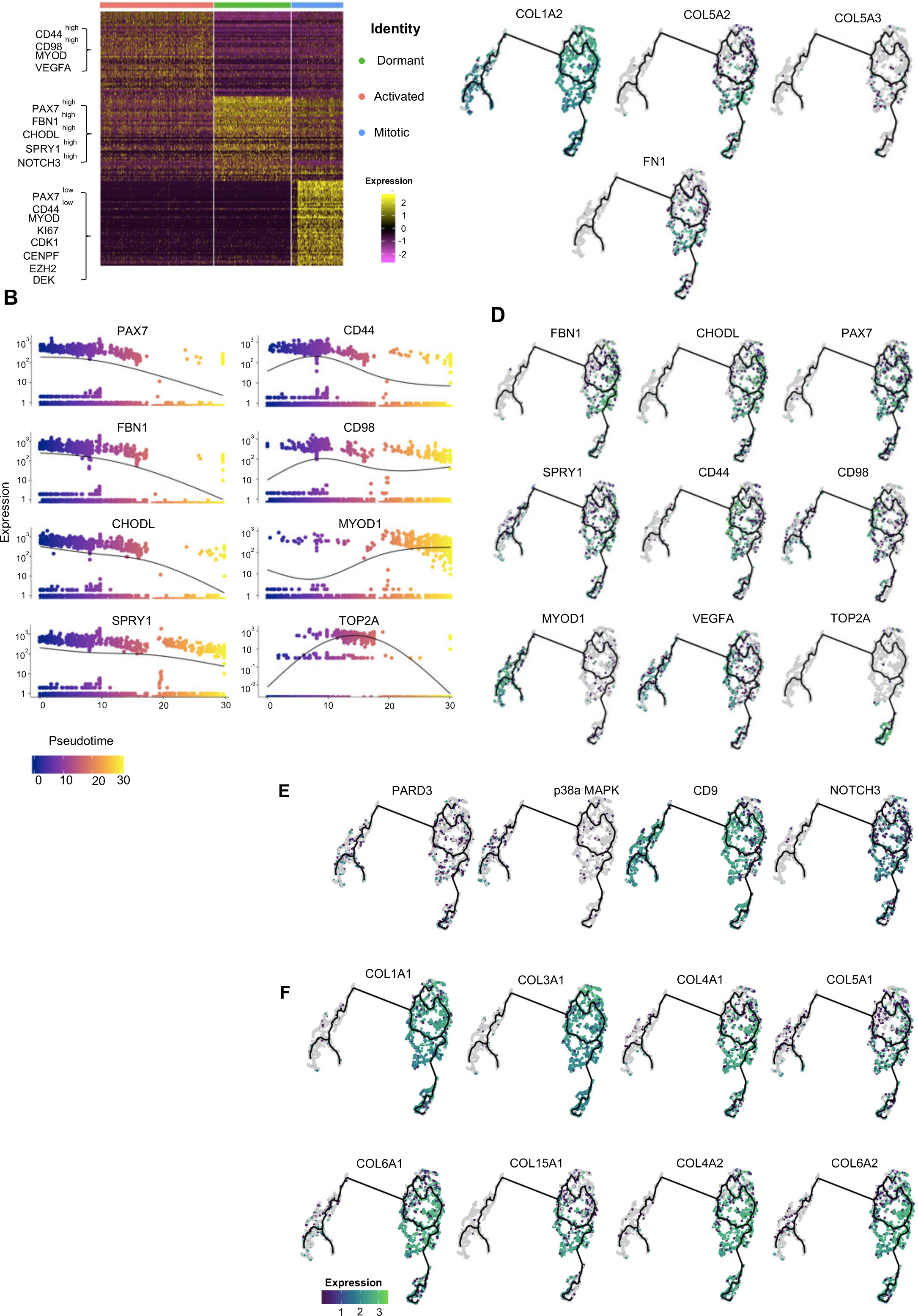
Pseudotime ordering of myogenic progenitor revealing distinct states and cell fate decisions. **(A)** Gene signatures of t-SNE described clusters based on relative expression levels of the 50 most significant markers for each of the 3 clusters **(B)** Expression of selected genes along pseudotime. Group of genes selected for myogenic progenitors: PAX7, SPRY1, CHODL (dormant state), CD44, CD98, MYOD1 (activated state), TOP2A (mitotic state), and for myoblasts: MYOD1 **(C, D)** UMAP feature plots depicting relative expression of extracellular matrix proteins COL1A2, COL5A2, COL5A3, FN1 (C) and selected transcriptional regulators that define dormant (PAX7, FBN1, CHODL, SPRY1), activated (CD44, CD98, MYOD1, VEGFA) and mitotic (TOP2A) state of myogenic progenitor and myoblast (MYOD1) clusters (D). **(E,F)** UMAP feature plots depicting relative expression of genes regulating asymmetric divisions and self-renewal (PARD3, p38a/b, CD9, NOTCH3) **(E)** and extracellular matrix collagens (COL1A1, COL3A1, COL4A1, COL5A1, COL6A1, COL15A1; COL4A2, COL6A2 (F).

**Figure 4 – figure supplement 3.**
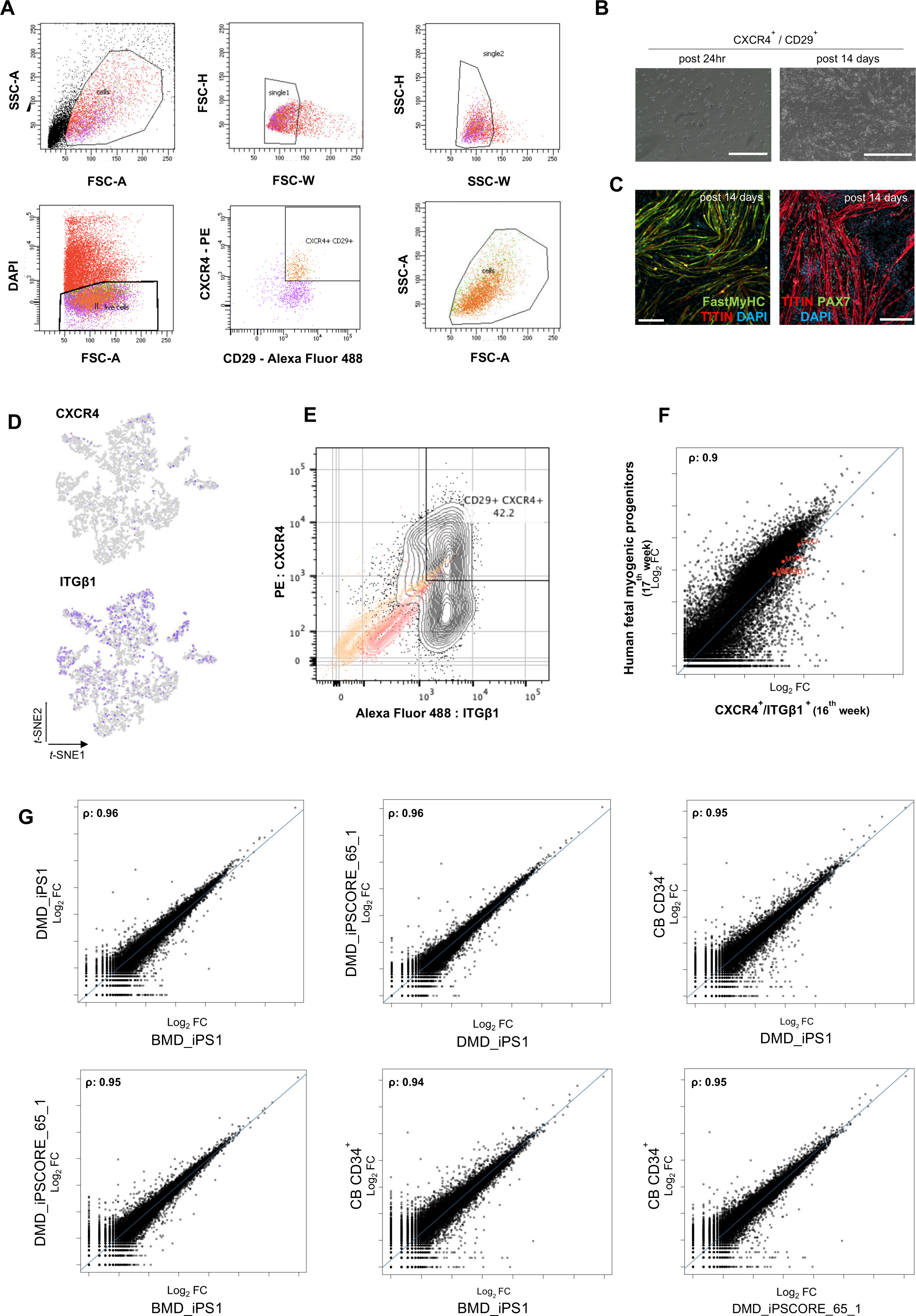
Organoid derived myogenic progenitors and correlation to fetal muscle progenitors. **(A)** Gating strategy applied for FACS sorting CD29+ / CXCR4+ cells from 15 -16 weeks skeletal muscle organoids **(B,C)** Re-plating CD29+ / CXCR4+ cells and culturing for 14 days highlights fetal myogenic potential, illustrated by brightfield and immunocytochemistry images for Fast MyHC+, TITIN+ and PAX7+ populations **(D)** t-SNE feature plots for ITGβ1 and CXCR4 demarcate expression of FACS isolated calls into myogenic progenitor subcluster **(E)** Gating strategy from (A) together with unstained population (yellow), isotype control (red) and CD29+ / CXCR4+ (gray) population **(F)** Correlation coefficient plot for Log2 fold change (Log2 FC) values for isolated myogenic progenitors from human fetal tissue (17w) and FACS sorted CXCR4+ / ITGβ1+ organoid derived myogenic progenitors (16w). PAX7, MYF5, MYOD1, MYOG are highlighted on the plot **(G)** Correlation coefficient plot for Log2 fold change (Log2 FC) values for FACS sorted CXCR4+ / ITGβ1+ organoid derived myogenic progenitors (16 w) from CB CD34+, DMD_IPS1, BMD_iPS1 and iPSCORE_65_1 pluripotent lines. Scale bars, 200uM (B), 100uM (C).

**Figure 4 – figure supplement 4.**
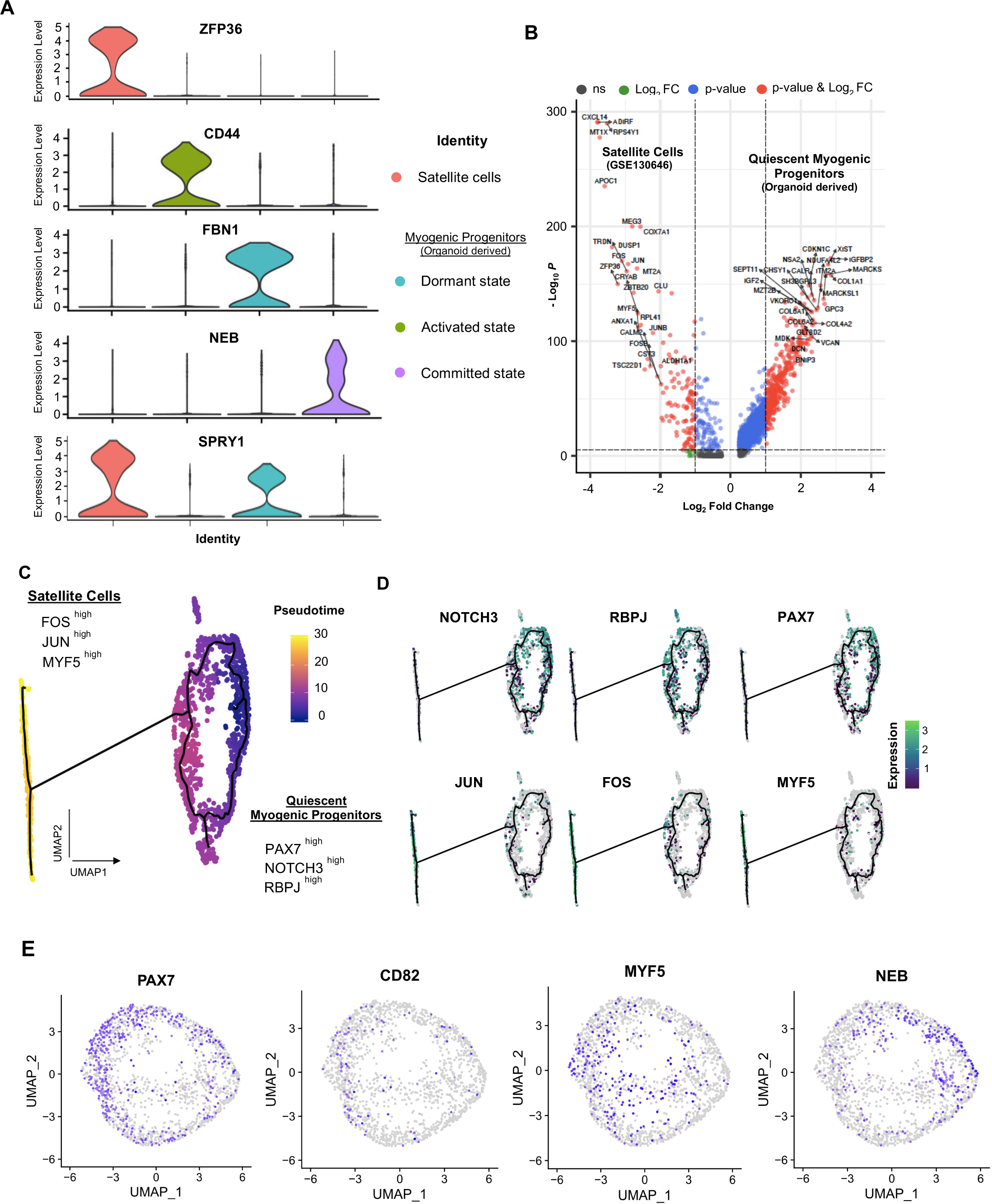
Organoid derived myogenic progenitors and correlation to adult human satellite cells. **(A)** Violin plots highlighting developmental status for organoid derived myogenic progenitors and satellite cells through relative expression of key signature markers ZFP36, CD44, FBN1, NEB and SPRY1 for each subcluster **(B)** Enhanced Volcano plot from comparing transcript levels between all adult satellite cells (GSE130646) and organoid derived myogenic progenitors. Log2 fold-change in normalized gene expression vs. - Log_10_ adjusted p-value is plotted. Differentially expressed genes: blue, adjusted p-value < 0.001; green, adjusted p-value > 0.001 and log2 fold-change > 1;,red, log2 fold-change > 1 and adjusted p value < 0.001; gray, no significance. **(C)** Pseudotime ordering for organoid derived myogenic progenitors and adult satellite cells together with expression of selected genes **(D)** Expression of genes selected for dormant myogenic progenitors (PAX7, NOTCH3, RBPJ) and activated satellite cells (MYF5, JUN, FOS). **(E)** UMAP feature plots depicting expression of PAX7, CD82+, MYF5 and NEB within the myogenic progenitor and adult satellite cell clusters corresponding to Figure 4F.

**Figure 4 – figure supplement 5.**
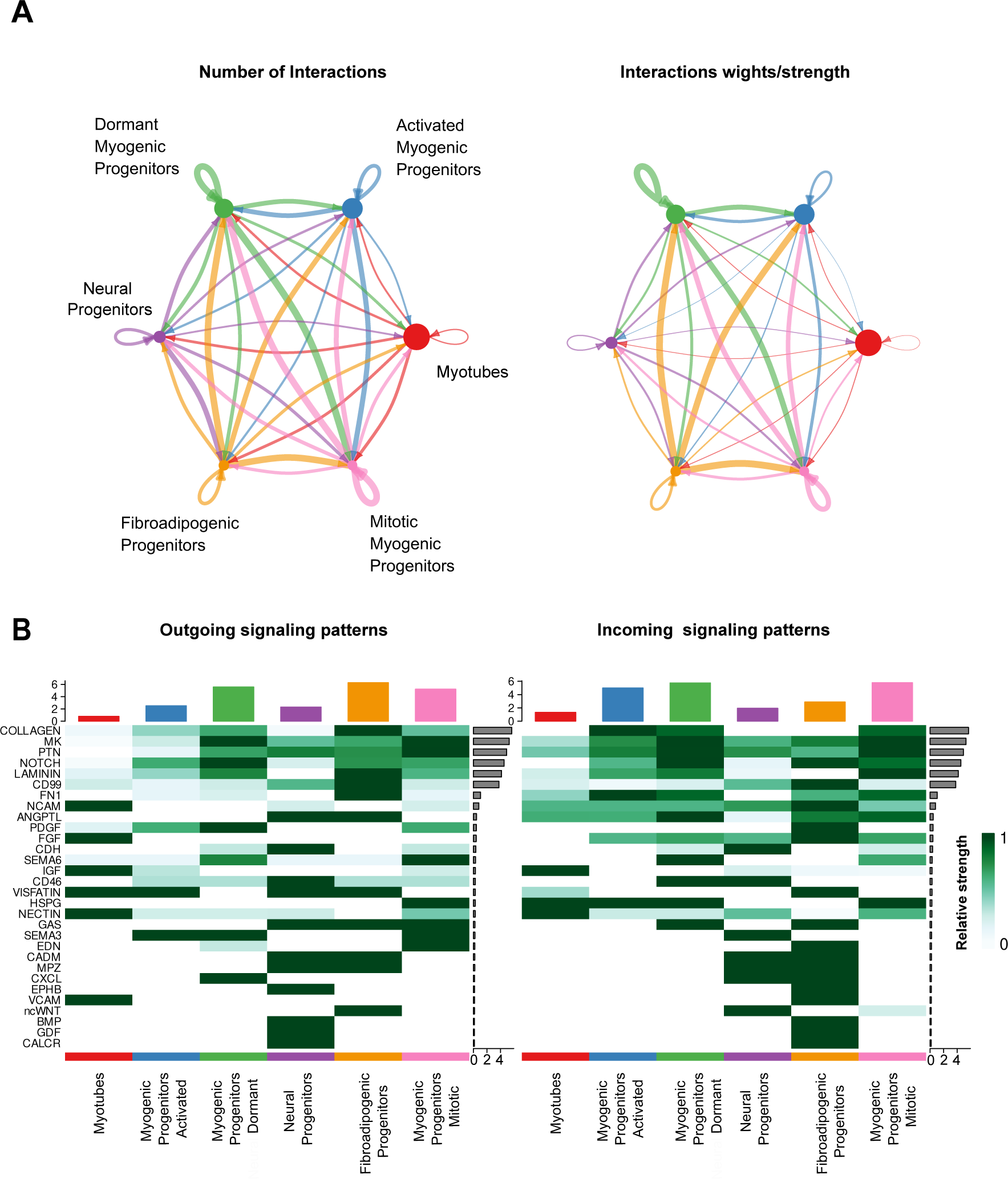
Characterization of cell-cell communication network for all clusters at week 12 of human skeletal muscle organoid development. **(A)** Circle plot illustrates the aggregated cell-cell communication network for all clusters at week 12 of human skeletal muscle organoids development. Circle sizes are proportional to the number of cells in each cell group and edge width represents the communication probability. **(B)** Heatmap indicating signals contributing most to the outgoing or incoming signalling among activated, mitotic, dormant/specification resistant myogenic progenitors and fibroadipogenic progenitors, neural progenitors and myofibers related clusters at week 12 of human skeletal muscle organoid development.

**Figure 4 – figure supplement 6.**
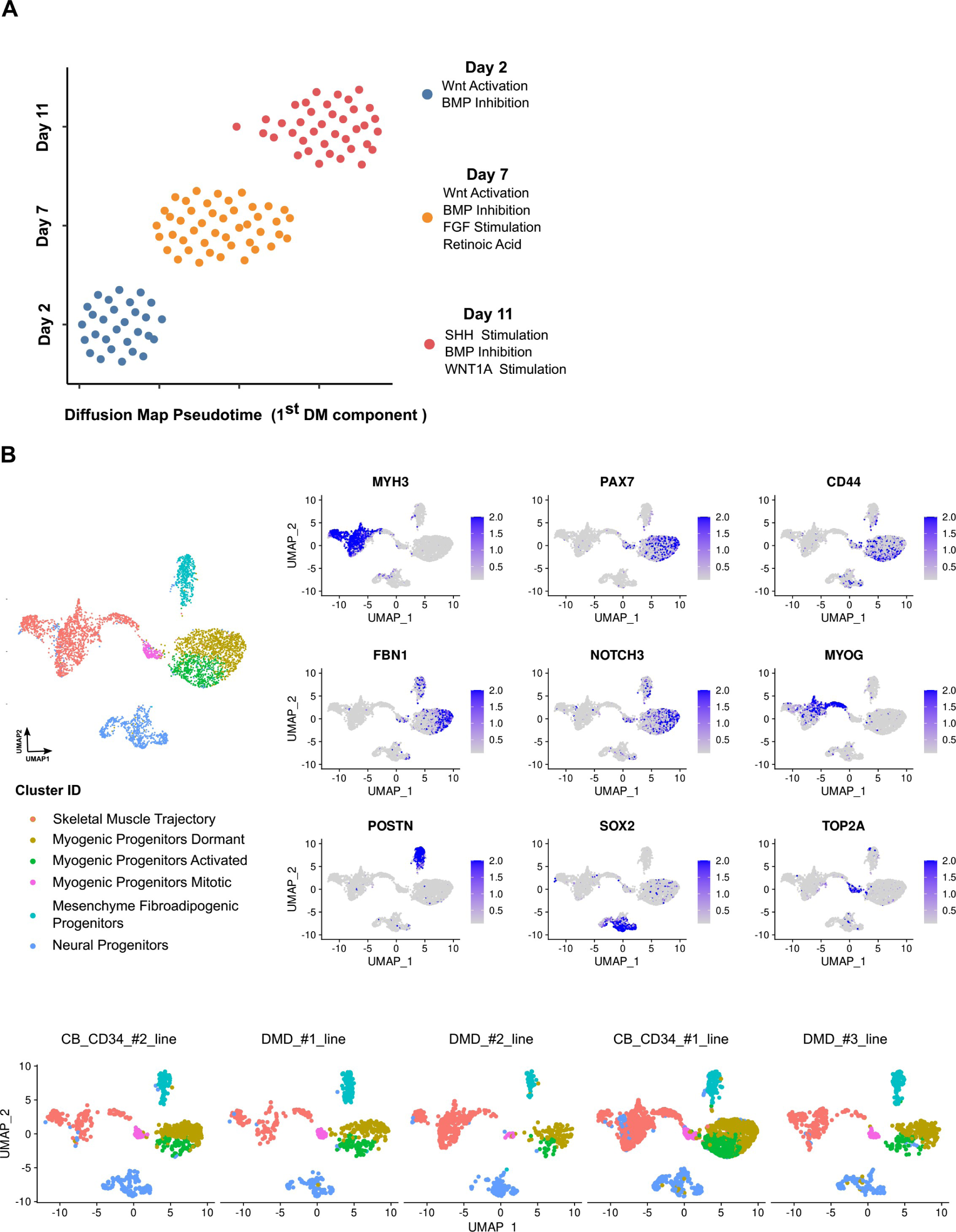
Reproducibility of organoid culture at early and mature stages. **(A)** Pseudotemporal ordering of organoids from Day 2, 7 and 11 based on qPCR expression profiling of selected 32 genes indicates robust homogeneous development of organoids during WNT activation and BMP Inhibition, while the subsequent development of paraxial mesoderm and neural tube related lineages via bFGF, Retinoic Acid (Day 7) and WNT1a, hSHH (Day 11) stimulation, introduced small variations to the organoid culture system. **(B)** Integrative analysis between datasets of organoids from different lines demonstrate highly conserved lineage representation for all datasets at mature stages (week 12) of human skeletal muscle organoid development. Feature plots on PAX7, CD44, FBN1, MYH3, POSTN, SOX2 as well as NOTCH3, MYOG, TOP2A indicate presence of dormant, activated and mitotic myogenic progenitors together with skeletal muscle myofibers, fibroadipogenic progenitors and neural progenitors related clusters.

**Figure 4 – figure supplement 7.**
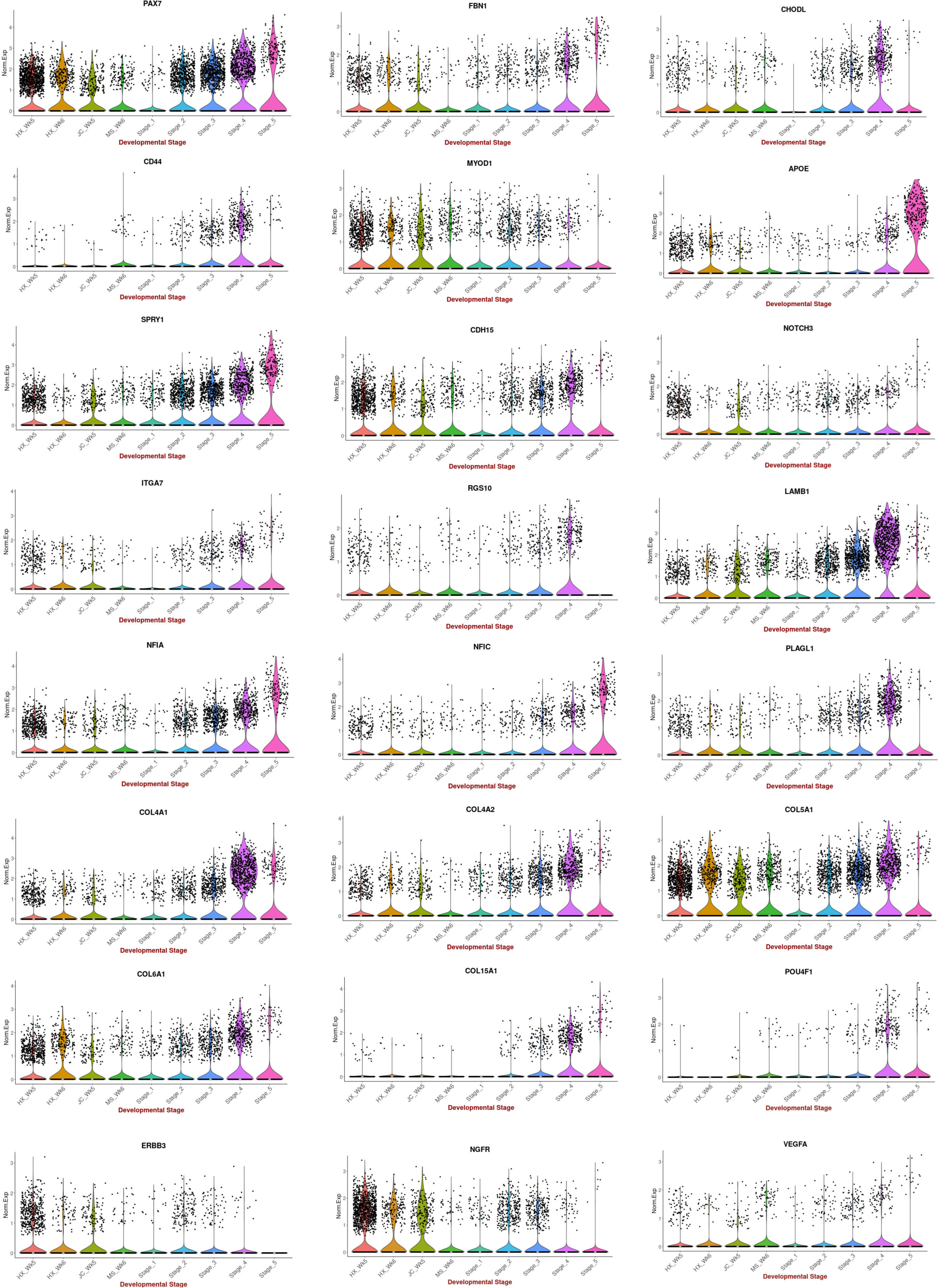
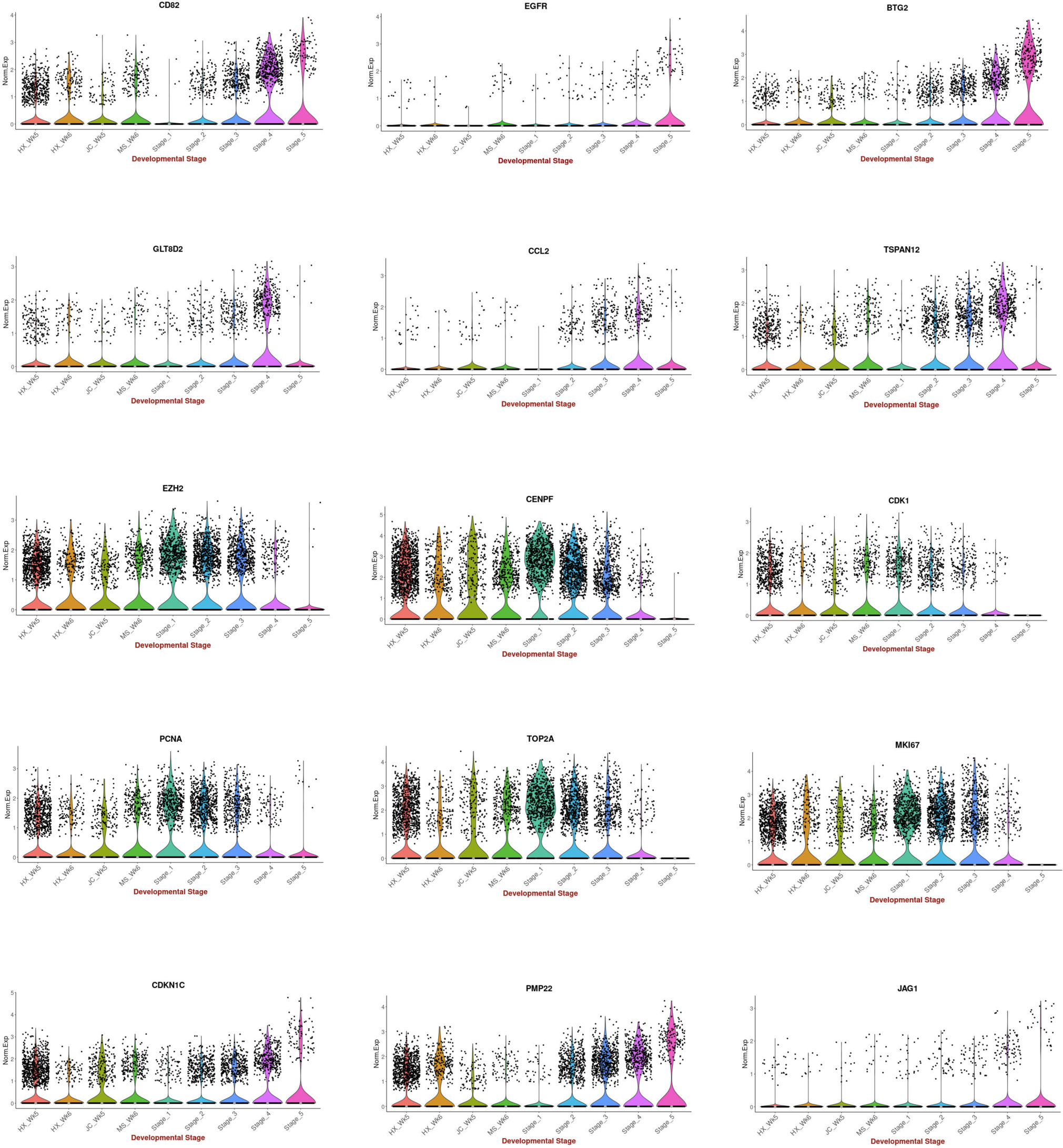
Comparison between 2D in vitro myogenic differentiation protocols and in vivo staging. Violin plots highlighting expression of key myogenesis markers: Comparison between myogenic progenitors derived in 2D differentiation protocols (HX protocol, Xi et al., 2017; JC protocol, Chal et al., 2015; MS protocol, Shelton et al., 2014) and in vivo myogenic progenitors (5 stages: 1,2: embryonic; 3,4: fetal; 5: postnatal). Data were acquired from Pyle ′s LAB UCLA website, (https://aprilpylelab.com/datasets/in-vivo-and-in-vitro-multiple-protocols-smpcs-and-scs/) and Xi et al. (2020).

**Table S1.**
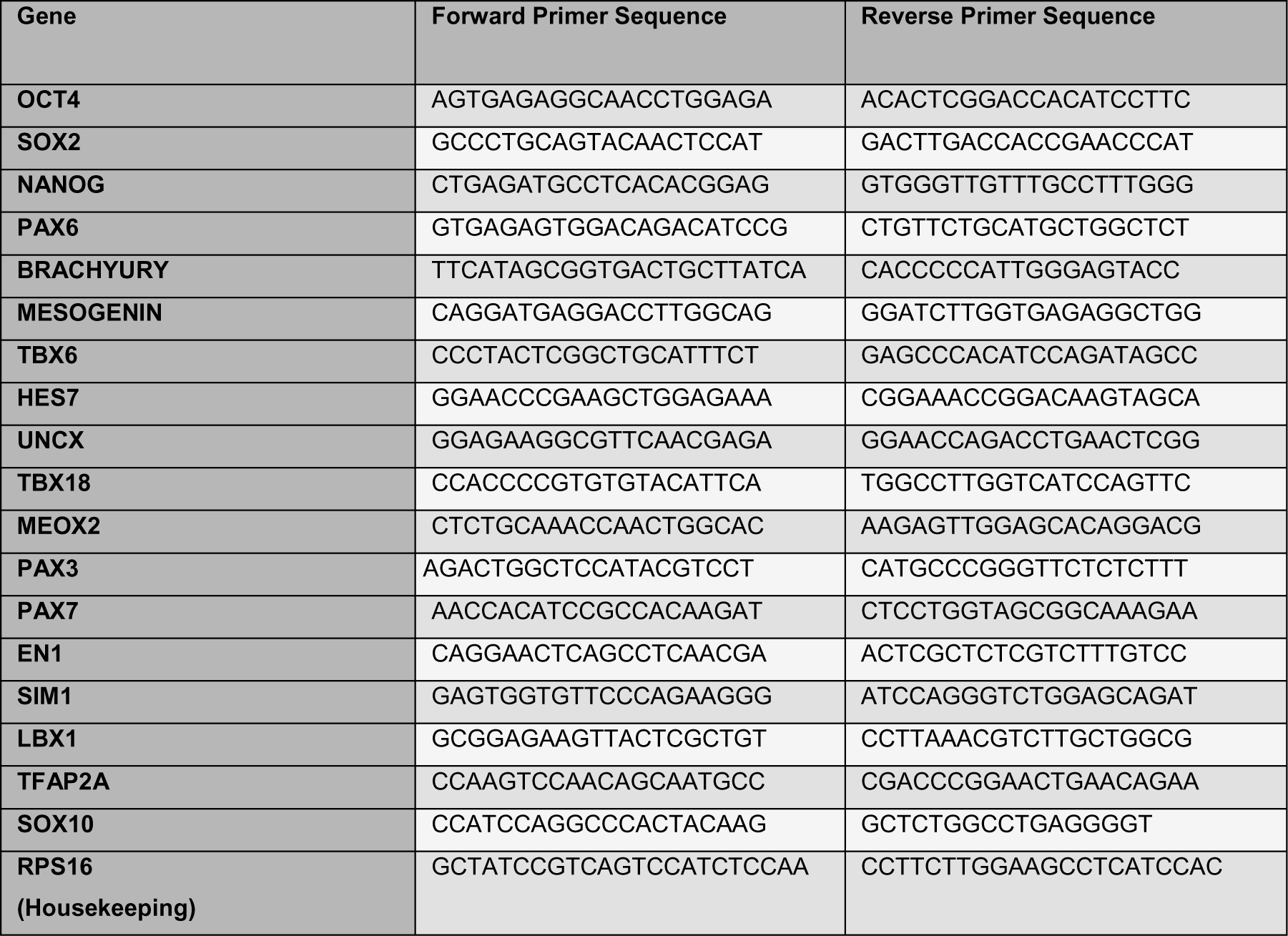
qPCR primer pairs applied to detect relative expression of key markers during skeletal muscle organoid development.

**Table S2.**
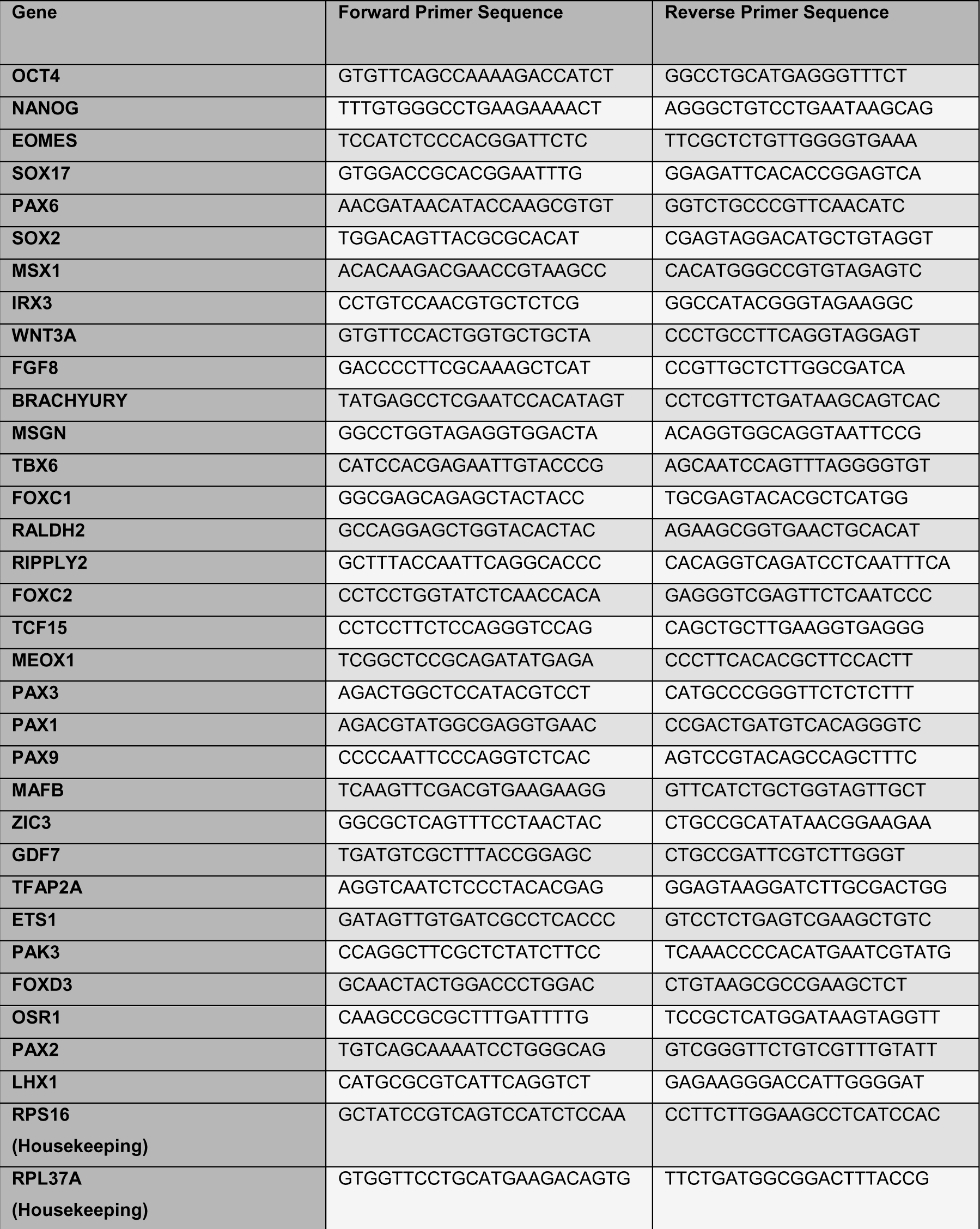
qPCR primer pairs applied for diffusion map analysis of early skeletal muscle organoid development.

